# SUMO2 Deletion Changes Chromatin Accessibility and Enhances Cytotoxic T Cell Activation and Tumor Infiltration

**DOI:** 10.64898/2026.01.27.700738

**Authors:** Mohottige D Neranjan Tharuka, Dai-Hua Chen, Maria Luisa Jurgensen Amaral, Tianchen Ren, Yixuan Kuang, Shih-Ting Huang, Nikhil Chilakapati, Bing Ren, Stephen P. Schoenberger, Ye Zheng, Yuan Chen

## Abstract

Cytotoxic T cells (CTL) are crucial for adaptive immunity that leads to prolonged survival and potential cures for cancer. Recent clinical data has shown that pharmacological inhibition of SUMOylation (SUMOi) profoundly modifies tumor microenvironment (TME) and activates CTL, although the mechanism is not well described. In this study, we found that T cell specific knock out (KO) of the most dominant SUMO paralog, *Sumo2*/*SUMO2*, in both mouse and human CD8^+^ T cells significantly enhanced CD8^+^ T cell activation that is independent of the known mechanism – inducing type I IFN (IFN-I) expression by myeloid cells. *Sumo2*/*SUMO2* KO in CD8^+^ T cells increased chromatin accessibility for transcription factors BATF, JunB, ATF3, FRA1, FRA2, and AP1 that are known to promote T cell activation and proliferation. Using antigen-specific T cell models, OT1 and Chimeric Antigen Receptor (CAR)-T cells, we found that *Sumo2* KO CD8^+^ T cells had significantly higher tumor infiltration as revealed by flow cytometry, immuno-fluorescence (IF) staining, and single nuclei RNA-sequencing (snRNA-seq) and conferred greater tumor growth inhibition than wildtype (WT) control T cells. snRNA-seq also revealed *Sumo2* KO CD8^+^ T cells increased the expression of Tumor Necrosis Factor-Related Apoptosis-inducing Ligand (TRAIL), induced apoptosis genes in tumor cells and activated IFN-I and IFN-γ responsive genes in all cell types in the TME. These findings elucidate a novel mechanism regarding how SUMOylation can directly control CTL activation and tumor infiltration that activate anti-tumor immunity in the TME. SUMO2 KO can also be a potential strategy to enhance adoptive T cell therapies of solid tumors by enhancing their activity, tumor infiltration and their ability to after the TME.

## Introduction

Cancer immunotherapy, including immune checkpoint inhibition and adoptive T cell therapy, has demonstrated the importance of inducing adaptive immunity in prolonged survival and cures for cancers. However, existing immune therapies have not been effective for most cancer types from hematological malignancy to solid tumors. Therefore, it is necessary to address novel immuno-oncology mechanisms to induce adaptive antitumor immunity to produce durable responses for more patients.

Inhibiting post-translation modifications by the small ubiquitin-like modifiers (SUMO) in dendritic cells and macrophages induces the expression of IFN-β (IFN-I) and IFN-γ target genes (*1*). The SUMO proteins are a subset of the ubiquitin-like proteins that can form covalent bonds with other proteins through a biochemical mechanism that is analogous to ubiquitination. SUMOylation requires several steps that are catalyzed by three enzymes; an activating enzyme E1 (which is a heterodimer of SAE1 and UBA2 (a.k.a. SAE2), a conjugation enzyme E2 (a.k.a. Ubc9) and one of ∼20 E3 ligases. The modifications can be removed by deconjugation enzymes to regulate the dynamic conjugation and deconjugation processes. SUMOylation does not directly lead to protein degradation, but adds a new docking site to target proteins, and thus enables protein complex formation through the SUMO-interacting motif (SIM) in receptor proteins (*2, 3*). Overexpression of the SUMO E1 and E2 enzymes is a hallmark of human cancers of many types, and the overexpression is a poor prognosis marker (*4-7*). The current prevailing view is that SUMOylation-induced IFN-I expression activates the observed anti-tumor immune effects (*1, 8*).

A phase I monotherapy clinical trial of the SUMO E1 inhibitor, TAK-981, has recently been completed (*9*). TAK-981 was well tolerated and showed profound tumor immune microenvironment (TME) modulation, including increased infiltration of CD8^+^ and CD4^+^ T cells, activation of CD4^+^ and CD8^+^ T cells in the TME, and increased expression of MHC-I on tumor cells in various solid tumors including melanoma at doses much lower than the maximum tolerated dose (MTD) (*9*). In addition, tumor control was observed including stable diseases for as long as three years and partial responses across different tumor types including melanoma, sarcoma, head and neck cancer, breast cancer, and pancreatic cancer, a proof-of-concept for the target (*9*).

In this study, we investigate whether suppressing SUMOylation in CTL can directly activate CTL independent of the IFN-I induction in dendritic cells and macrophages. Using antigen-specific CD8^+^ T cell models, OT1 and Chimeric Antigen Receptor (CAR)-T cells, we found that T cell specific knock out (KO) of the most dominant SUMO paralog, *Sumo2*/*SUMO2*, in human and mouse CD8^+^ T cells significantly enhanced CD8^+^ T cell activation, effector function and tumor infiltration. *Sumo2*/*SUMO2* KO of CD8^+^ T cells increased chromatin accessibility for transcription factors known to promote T cell activation and proliferation, including BATF, JunB, ATF3, FRA1, FRA2, and AP1. Notably, *Sumo2* KO CD8^+^ T cells have increased infiltration in the TME compared to the wildtype (WT) littermate control T cells as revealed by flow cytometry, immuno-fluorescence (IF) staining, and single nuclei RNA sequencing (snRNA-seq). Although *Sumo2* KO in mouse CD8^+^ T cells had reduced expression of type I IFN responsive genes upon *in vitro* activation, snRNA-seq revealed that these CD8^+^ T cells induced significant IFN-γ and IFN-I target genes in all cell types in the TME, including myeloid cells and tumor cells. Analysis of cell-cell interactions in the TME using snRNA-seq data revealed that T cell-specific *Sumo2* KO increased the expression of Tumor Necrosis Factor-Related Apoptosis-inducing Ligand (TRAIL) on CD8^+^ T cells and activation of apoptotic genes in tumor cells. The findings elucidate how the post-translational modification with SUMO can directly control CD8^+^ T cell activation, effector function, tumor infiltration and tumor inhibition. The mechanistic insights revealed in our studies will be informative for designing new T cell therapy and for designing effective combination immune therapy incorporating SUMOylation suppression as a new strategy.

## Results

### KO of *SUMO2/Sumo2* in both human and mice T cells enhanced their activation

SUMO2 is the most dominant SUMO paralog in both humans and mice T cells (**Fig. 1A**). The mRNA levels of each SUMO paralogs were measured by reverse transcription-quantitative polymerase chain reaction (RT-qPCR). In mice, *Sumo2* is the dominant isoform accounting for >65% of the total *Sumo* gene expression in T cells of adult mice, while *Sumo1* and *Sumo3* account for ∼25% and ∼9% respectively of total *Sumo* expression (**Fig. 1A**). This finding is consistent with a previous report that *Sumo2* expression is higher than other *Sumo* paralogs in adult heart, brain and kidney tissues (*10*). In human CD8^+^ T cells, *SUMO2* is also the dominant paralog, accounting for approximately ∼76% of the total *SUMO* gene expression. *SUMO1* and *SUMO3* account for 11% each. Human cells express an additional *SUMO4*, which is the least expressed *SUMO* gene, accounting for less than 1% of the total *SUMO* gene expression (**Fig. 1A**).

**Figure 1.**
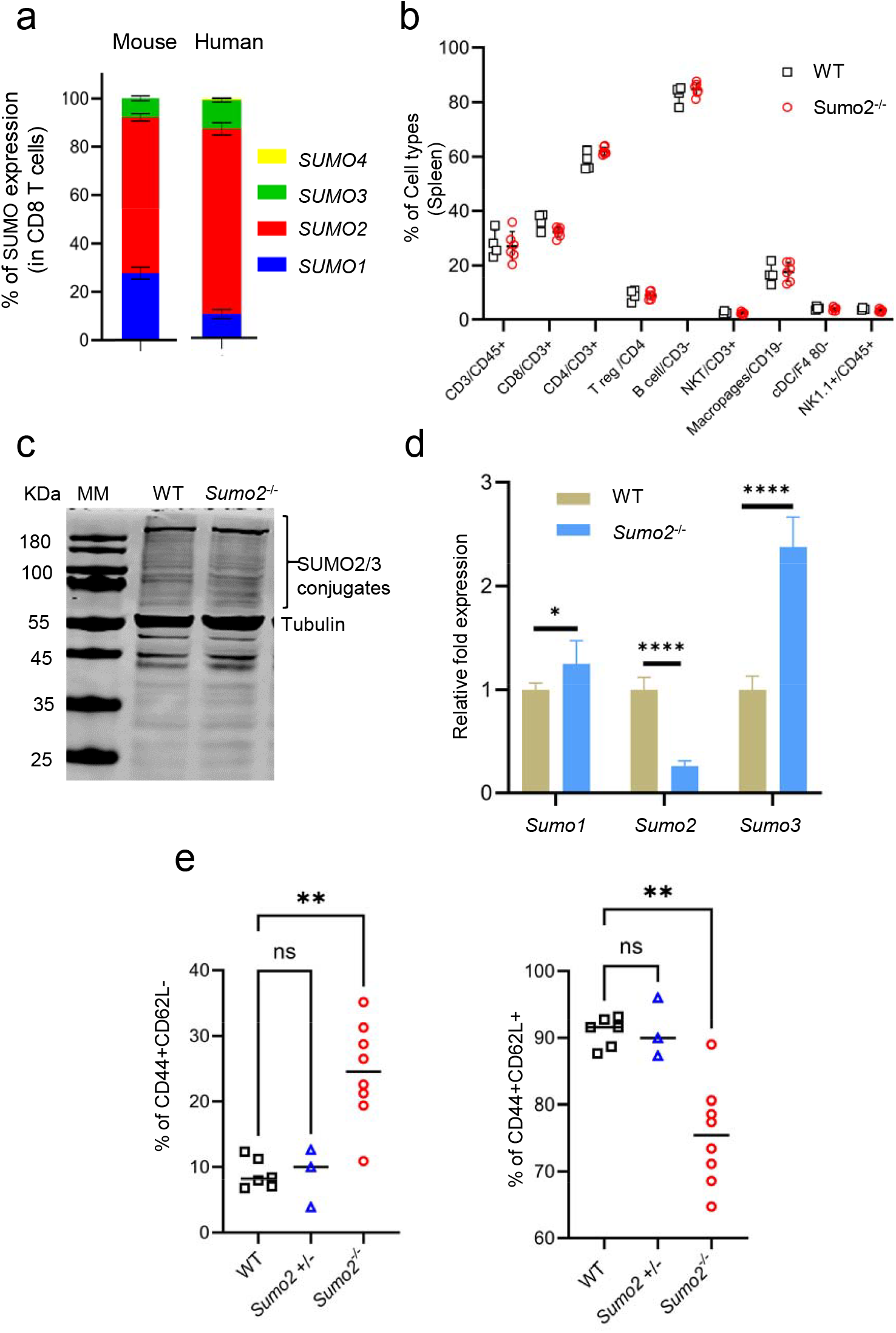
Knocking out the most abundant SUMO2 isoform in mice enhances effector CD8^+^ T cell phenotypes without altering other immune cell populations. **(A)** RT-qPCR analysis of Sumo/SUMO isoforms in Human and Mouse CD8+ T cells. Results show mean and SD of three independent experiments (n=3). **(B)** Flow cytometric analysis of splenocytes from WT and *Sumo2*^−/−^ mice showing the major adaptive immune cell populations (CD3^+^, CD8^+^, CD4^+^, and Tregs) or innate immune cell subsets (NKT cells, NK cells, macrophages, and cDCs) (n=3). **(C)** Western blot analysis of total SUMO2/3 conjugate protein levels between WT and *Sumo2*^−/−^ mouse activated T cells. The current SUMO2/3 antibodies do distinguish between SUMO2 and SUMO3 isoforms. Tubulin antibody was used as reference control (n=2). **(D)** RT-qPCR analysis of *Sumo1,Sumo2* and *Sumo3* isoforms expression in *Sumo2*^−/−^ mouse T cells. Mouse GAPDH was used as reference control (WT: n=3; *Sumo2*^−/−^ n=5). **(E)** Flow cytometric analysis of WT, *Sumo2*^+/−^, *Sumo2*^−/−^, CD8^+^ T cells upon activation. CD44^+^CD62L^−^ effector cells (EF) and CD44^+^CD62L^+^ central memory-like cells (CM) indicated as mean and SD (WT: n=6; *Sumo2*^+/−^ n=3; *Sumo2*^−/−^ : n=8).

Because *Sumo2*/*SUMO2* is the dominant *SUMO* paralog, we created *Sumo2* floxed mice and obtained T cell specific deletion of *Sumo2* by breeding with Lck-*Cre* transgenic mice to delete *Sumo2* systemically in T cells (**Fig. S1**). *Sumo2* KO mice have no abnormal phenotype. This contrasts with the fact that systemic *Sumo2* deletion is embryonic lethal (*10*), suggesting that T cell specific *Sumo2* KO does not affect development. In addition, *Sumo2* KO does not affect circulating immune cell populations (**Fig. 1B**).

Western blots of T cells show that global SUMOylation is only slightly reduced (**Fig. 1C**). This is due to increased expression of *Sumo3*, which is nearly identical *Sumo2* in amino acid sequence and there is no antibody to distinguish between the two. RT-qPCR results clearly demonstrate that SUMO2 expression is reduced by approximately 75%, whereas SUMO3 expression is elevated in T cells. (**Fig. 1D**). However, *Sumo2* homozygous KO T cells showed increased population of the effector phenotype and reduced population of central memory phenotype (**Fig. 1E, Fig. S2**). In contrast, *Sumo2* heterozygous deletion in CD8 T cells produced similar effector and central memory populations as the WT T cells (**Fig. 1E, Fig. S2**).

T cell activation is critical to T cell effector function and differentiation. To determine how *Sumo2* KO affects T cell activation, CD8^+^ T cells were isolated from the splenocytes of *Sumo2* KO mice and littermate control mice (WT). Upon T cell receptor (TCR) stimulation using anti-CD3/CD28 antibodies, markers for T cell activation and exhaustion were analyzed by flow cytometry at different time points after TCR stimulation. *Sumo2* KO increased CD8^+^ T cell activation as shown by increased populations of CD69, IFNγ, TNFα, IL2 and CD25 expression cells (**Fig. 2A-2E**). In addition, the costimulatory receptor OX40 expression T cells also increased (**Fig. 2F**). PD1 expression CD8^+^ T cells increased by TCR stimulation that may reflect activation (**Fig. 2G**).

**Figure 2.**
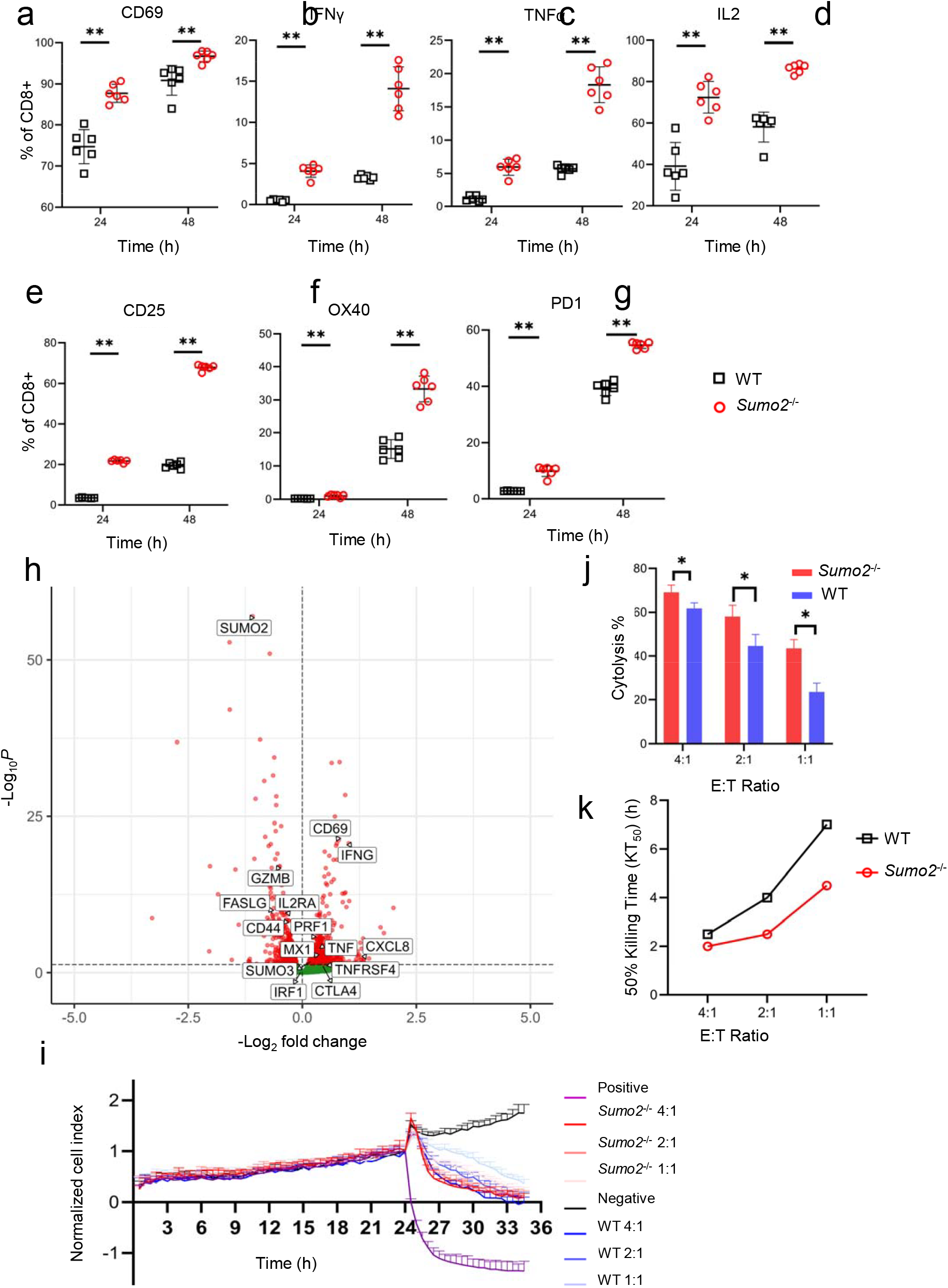
SUMO2 deficiency enhances CD8^+^ T cell activation, effector function, and cytotoxic capacity. **(A–G)** Flow-cytometric analysis of mouse CD8^+^ T cells activated for 24 h or 48 h. The expression of activation and effector markers determined include CD69, IFN-γ, TNF-α, IL-2, CD25, OX40, and PD-1 at both time points in *Sumo2*^−/−^ compared with WT controls (n=6). **(H)** Volcano plot of differentially expressed genes analyzed with DESeq2 from bulk RNAseq data of *SUMO2*^*-*/-^ (left) or WT (right) human CD8+ T cells. Statistically significant (p-value < 0.05) genes are highlighted in red and not significant genes are highlighted in green. **(I)** Real-time cytotoxicity assay using an xCELLigence analyzer. Mouse *Sumo2*^−/−^ and WT CD8^+^ CAR T cells were co-cultured with antigen-specific B16F0-human-CD19 melanoma cells across multiple effector-to-target (E:T) ratios (n=3). **(J)** Cytolysis calculation at 3 h post-co-culture of *Sumo2*^−/−^ and WT CAR-T cells with antigen-specific B16F0-hCD19 melanoma cells at 4:1, 2:1, and 1:1 E:T ratios (n=3). **(K)** KT_50_ values, representing the time required to kill 50% of target cells, *Sumo2*^−/−^ CAR T cells compared with WT CAR T cells across all E:T ratios, indicating accelerated tumor-cell killing. Statistical analysis: Mann–Whitney tests (panel A-E) and t-test (G-H) \**P* < 0.05, \*\**P* < 0.01.

To determine whether the enhanced CD8^+^ T cell activation upon *Sumo2* KO occurs in human T cells, human CD8^+^ T cells were isolated from PBMC (Peripheral Blood Mononuclear Cells) of healthy donors. Following TCR stimulation of the human CD8^+^ T cells by Dynabeads™ Human T-Activator CD3/CD28 and cultured with the addition of IL2, CRISPR was used to achieve approximately 80% *SUMO2* KO as shown by qPCR (**Fig. S3**). *SUMO2* KO also increased the expression of *SUMO3* (**Fig. S3**), consistent with that observed in mice (**Fig. 1D**). RNA-seq data confirmed reduced expression of *SUMO2* **(Fig. 2H)**. In the human RNA-seq data, *SUMO2* deletion resulted in an increase in early activation marker *CD69* as well as increased expression cytokines related to effector function, including *IFNG, PRF1* and *TNF*, supporting that the enhanced activation observed in mouse CD8^+^ T cells is relevant to the human T cells (**Fig. 2H**).

To determine antigen-specific T cell activity, we used a mouse CAR-T model against engineered mouse tumor cells (*11*), in which a mouse tumor cell line, B16F0, was transduced to express human CD19 (hCD19) (B16F0-hCD19). Expressing hCD19 in mouse tumor cell lines did not change growth rates in syngeneic mouse models (*11*). A retroviral vector was used to transduce mouse T cells, wild-type (WT) or *Sumo2* KO, with an anti-hCD19 CAR (*11*). CAR expression in WT and *Sumo2*^-/-^ CD8^+^ T cells were determined to be over 90% **(Fig. S4)**. The CD19-CAR also expresses a congenic marker. Killing of B16F0-hCD19 cells by the *Sumo2* KO CD19-CAR T cells and WT control was measured with an xCELLigence Real Time Cell Analysis instrument (**Fig. 2I, Fig. S5**). *Sumo2* KO significantly enhanced CAR-T cell killing of tumor cells, as shown by the analysis of the percentage of cancer cells killed at 3 hours (h) (**Fig. 2J**). In addition, the faster killing of tumor cells by *Sumo2* KO mice is shown by the time it took to kill 50% of tumor cells (**Fig. 2K**). This is consistent with increased CD8^+^ T cell activation upon *Sumo2* KO.

### *Sumo2/SUMO2* KO increases chromatin accessibility for transcription factors important for T cell activation and proliferation

To understand the molecular mechanism of *Sumo2/SUMO2* KO induced CD8^+^ activation and effector function increase, we conducted genome-wide analysis using ATAC-sequencing (ATAC-seq) in both mouse and human CD8+ T cells. *Sumo2*/SUMO2 KO resulted in significantly more accessibility across the genome than control WT for both mouse and human CD8^+^ T cells, with 5152 peaks enriched in *Sumo2* KO vs. 932 peaks enriched in WT control in mouse CD8^+^ T cells, and 5727 peaks enriched in SUMO2 KO vs. 163 peaks enriched in WT control in human CD8^+^ T cells (**Fig. 3A**). In both mouse and human CD8^+^ T cells, HOMER motif analysis on peaks specifically enriched in either *Sumo2*/SUMO2 KO or WT control cells revealed different enriched motifs (**Fig. 3B**). The *Sumo2/SUMO2* KO CD8^+^ T cell-specific ATAC-seq peaks showed a strong enrichment of BATF, JunB, ATF3, FRA1, FRA2, and AP1, whereas control WT cell-specific peaks demonstrated moderate enrichment of ETS motifs (**Fig 3B**).

**Fig. 3.**
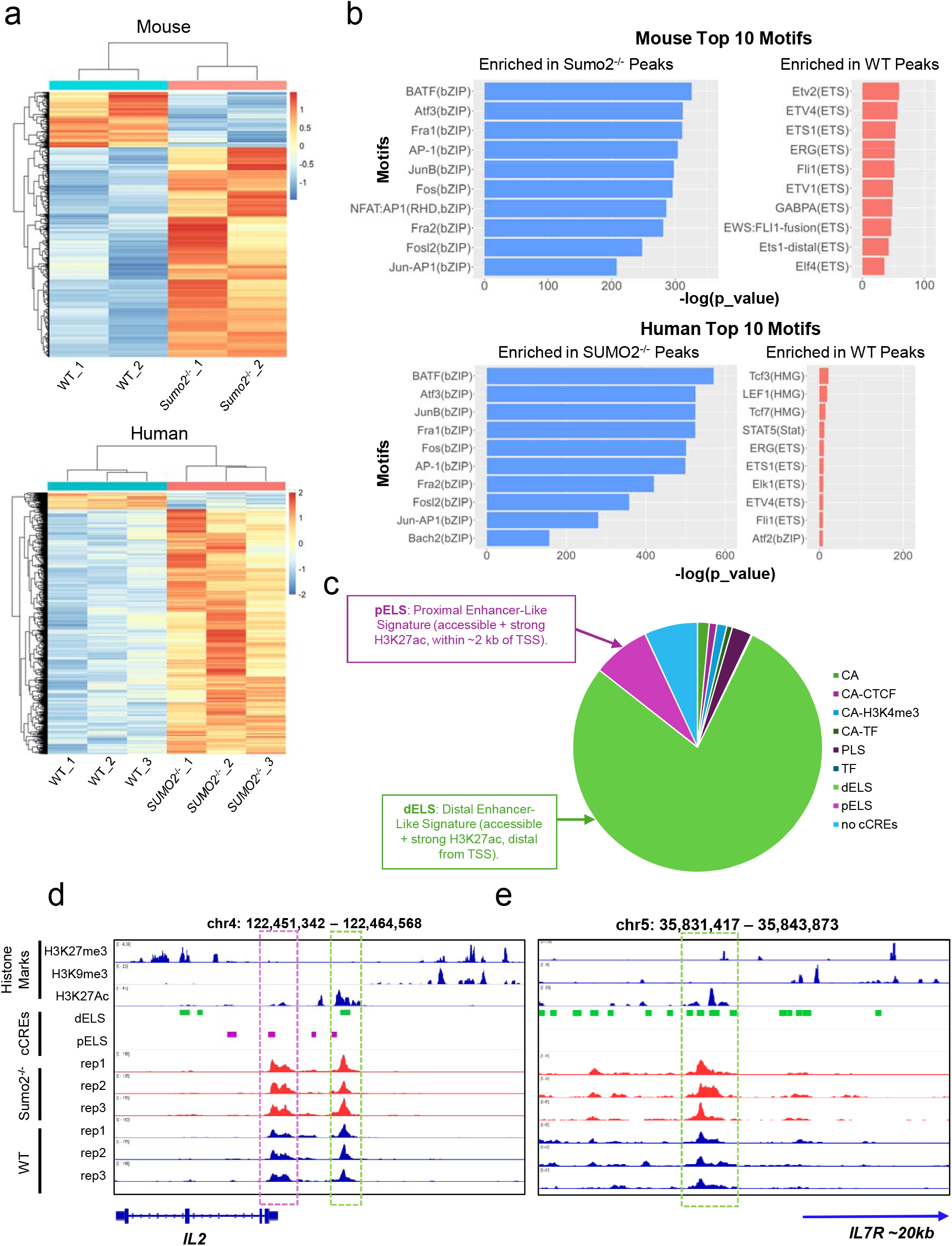

We analyzed association of the *SUMO2* knockout enriched peaks with annotated histone marks, in particular H3K27Ac, in human CD8^+^ T cells in the ENCODE database. We found that a significant number of peaks (1776 peaks) enriched in the SUMO2^-/-^ T cells coincided with known H3K27Ac marks **(Fig. S6)**. Because H3K27Ac marks are often associated with active enhancer regions in the genome, we annotated our *SUMO2*^-/-^ enriched peaks with candidate cis-Regulatory Elements (cCREs) annotations in the ENCODE database. We found that around 78% of all *SUMO2* knockout enriched peaks overlapped with known distal enhancer-like signatures (dELS), and around 7.8% overlapped with proximal enhancer-like signatures (pELS) **(Fig. 3C)**. Representative regions showing increase chromatin accessibility in pELS **(Fig. 3D)** and dELS regions **(Fig. 3E)** are overlaid with histone marks near the IL2 coding region and upstream of the IL7R coding region.

We also conducted Transcription Factor (TF) footprint analysis with TOBIAS using the CORE Vertebrates motifs from the JASPAR database. The TF footprint analysis corrects for transposon bias and uses patterns from the ATAC-seq signals to predict TF binding. For both humans and mice, we observe the same motifs as shown in ATAC-seq being exclusively enriched in *Sumo2* KO samples (**Fig. S7**, upper panel), consistent with our HOMER analysis. The footprint score of the top scoring motifs from the *Sumo2* knockout cells and control WT cells were then evaluated. Similar to the HOMER motif analysis, the motifs enriched in the *Sumo2* KO cells were very significant and differentially enriched in the *Sumo2* KO cells compared to the control cells, as indicated by the box plot (**Fig. S7**, lower panel). In contrast, TF footprint identification in WT control cells was not statistically significant (**Fig. S7**, lower panel). Taken together, these findings indicate that SUMO2 regulates chromatin accessibility of CD8^+^ T cells for transcriptional programs required for activation and proliferation.

### *Sumo2* KO increased proliferation, tumor infiltration and tumor control *in vivo*

We performed *in vivo* experiments to examine how the increased effector function of CD8+ T cells by Sumo2 KO influence *tumor growth*. The same CAR-T cells used for the *in vitro* studies (**Fig. 2**) were used for *in vivo* studies by implanting *Rag1*^-/-^ mice with B16F0-hCD19 tumors. On day 0, 8–10-week-old *Rag1*^-/-^ mice were injected subcutaneously with 1 × 10^5^ B16-hCD19 cells (**Fig. 4A**). On Day 6, 3×10^6^ CAR-transduced CD8^+^ T cells were adoptively transferred into tumor size-matched, tumor-bearing mice. Once tumors became palpable, tumor size was measured with a caliper every other day, and tumor volume was calculated as 0.5 × (length × width^2^). On Day 20, tumors, spleens, and blood were harvested for immune profiling of the TME by flow cytometry (representative data shown in **Fig. 4B**). Flow cytometry analysis revealed more *Sumo2* KO than WT CAR-T cells in the TME (**Fig. 4C**). In addition, *Sumo2* KO CAR-T cells have an increased trend in the spleen (**Fig. 4D**) and significantly increased in the PBMC (**Fig. 4E**), suggesting that there was an increase in CAR-T cell proliferation. This is consistent with BATF motifs being enriched in differentially accessible chromatin regions upon Sumo2 KO. Previous studies have shown that BATF is a TF important for T cell proliferation and activation (*12-14*). *Sumo2* KO CAR-T cells suppressed tumor growth more significantly than the WT CAR-T cells (**Fig. 4F**).

**Fig. 4.**
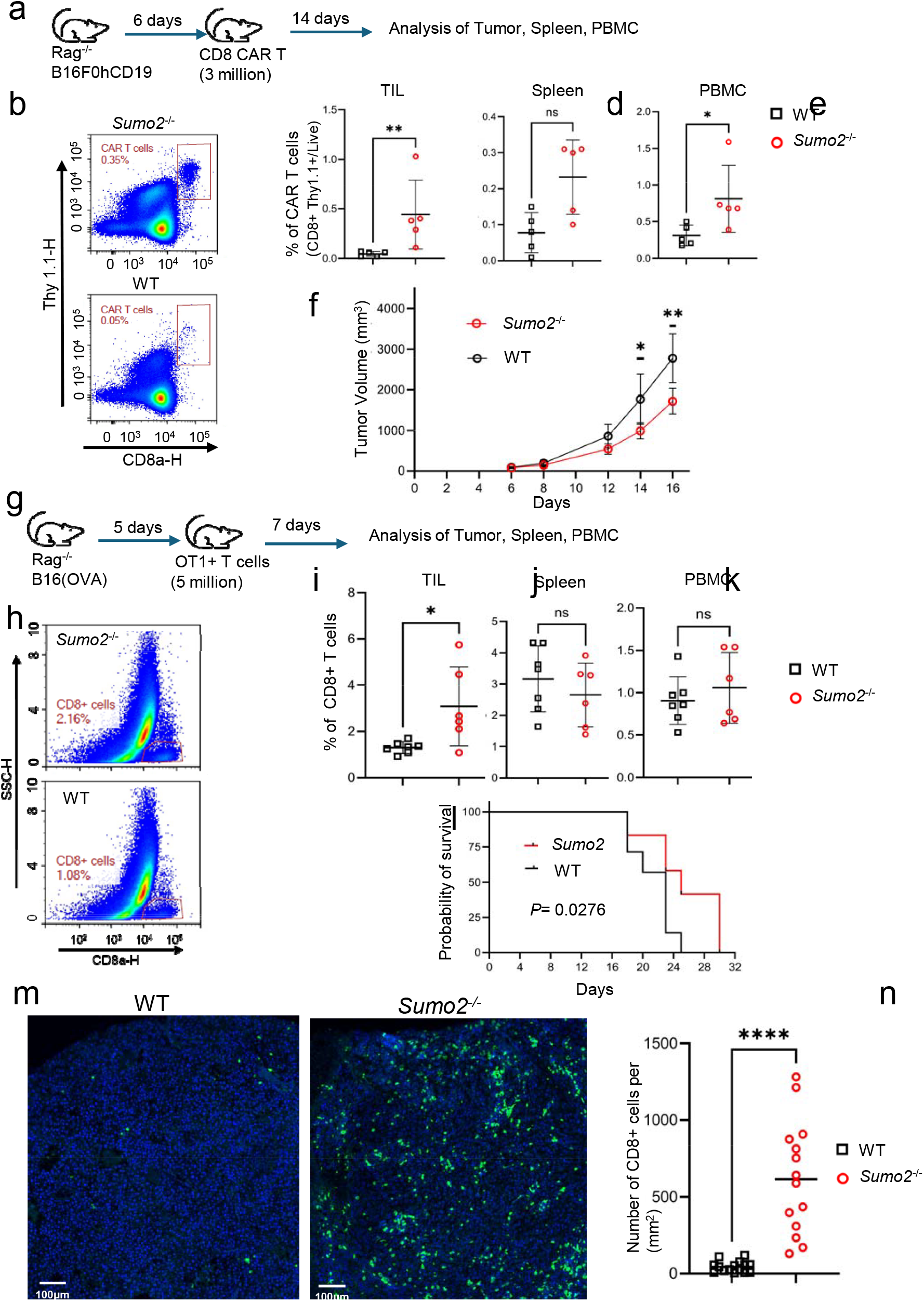
*Sumo2* KO enhances CAR-T cell and OT1 cell expansion, tumor infiltration, and tumor control *in vivo*. **(A)** Schematic of the CAR T-cell adoptive transfer to *Rag1*^−/−^ mice bearing tumors. B16F0-hCD19 tumor cells (1×10^5^) were injected subcutaneously on day 0. On day 6, 3×10^6^ CAR T cells were adoptively transferred via retro-orbital injection. Fourteen days after transfer, tissues (spleen, PBMCs, and tumor) were collected for flow cytometric analysis. **(B)** Representative flow cytometry plots showing CAR T cells (Thy1.1+, CD8+) within the tumor microenvironment (TME). **(C–E)** Quantification of CAR T-cell frequencies by flow cytometry in Tumor Infiltrating Lymphocytes (TILs), spleen, and PBMCs (n=5). **(F)** Tumor volume measurements in the CAR T-cell adoptive transfer model *in vivo*. *Sumo2*^−/−^ CAR T–treated mice has reduced tumor growth compared to WT controls (n=5). **(G)** Schematic of the experiment by OT1 adoptive transfer to *Rag1*^−/−^ bearing tumors. B16-OVA tumor cells (1×10^5^) were injected subcutaneously on day 0. On day 5, 5×10^6^ activated OT1 CD8+ T cells were transferred via retro-orbital injection. Tissues (spleen, PBMCs, and tumor) were harvested 7 days post-transfer for flow analysis. **(H)** Representative flow cytometry plots of total CD8^+^ T cells in the TME. **(I–K)** Frequencies of OT1 CD8^+^ T cells in TILs, spleen, and PBMCs (WT n=7; SUMO2^−/−^ n=6). **(L)** Survival analysis of the OT1 *Rag1*^−/−^ adoptive transfer model. Mice received B16-OVA tumor implantation on day 0 and activated OT1 T cells transfer on day 5 (WT n=7; SUMO2^−/−^ n=12; *P* = 0.0276). **(M–N)** Immunofluorescence staining of tissue sections from B16-OVA tumors following *Sumo2*^−/−^ and WT adoptive OT1 transfer. **(M)** Representative images demonstrating infiltration of OT1 CD8^+^ T cells (Green: anti-CD8, Blue: DAPI). **(N)** Quantification of intratumoral CD8^+^ T cells between the *Sumo2*^−/−^ group and WT group. Statistical analysis: Multiple Mann–Whitney tests (panel C-E,I-K), Logrank (Mantel-Cox Test) for survival (L), and unpaired T test for (N) \**P* < 0.05, \*\**P* < 0.01, \*\*\**P* < 0.001, and \*\*\*\**P* < 0.0001.

To determine the effects with endogenous T cell receptor (TCR), we conducted a similar experiment using *Sumo2* KO OT1 cells by crossing OT1 mice with *Sumo2* KO mice (**Fig. 4G**) and used WT OT1 as control. B16 cells expressing OVA antigen (B16(OVA)) were used to establish a tumor model in 8–10-week-old *Rag1*^-/-^ mice by subcutaneously injecting 1 × 10^5^ B16(OVA) cells. On Day 5, 5 × 10^6^ OT1 CD8^+^ T cells were adoptively transferred into tumor size-matched, tumor-bearing mice. On Day 12, tumors, spleens, and blood were harvested for immune profiling of the TME by flow cytometry (representative data shown in **Fig. 4H**). There were significantly more *Sumo2* KO than WT OT1 cells in the TME (**Fig. 4H, I**). However, OT1 cells in the spleen and PBMC were similar between the two groups (**Fig. 4J, K**). No differences in exhaustion markers were identified (**Fig. S8**). To determine tumor control, a survival experiment was conducted where mice were monitored until the experimental endpoint. *Sumo2* KO OT1 significantly prolonged survival compared to WT OT1 (**Fig. 4L**). Furthermore, IF staining of tumor slides more clearly demonstrated increased infiltration of *Sumo2* KO OT1 cells in the tumor tissue than WT OT1 cells than flow cytometry **(Fig. 4M, N)**.

Taken together, both models have shown that more *Sumo2* KO CD8^+^ T cells infiltrated into the tumors than WT CD8^+^ T cells. The increased CD8^+^ T cell tumor infiltration may, in part, be due to increased proliferation of the *Sumo2* KO T cells. *Sumo2* KO CD8^+^ T cells increase tumor control in both models, measured by tumor growth or survival.

### snRNA-seq reveals profound TME modulation by *Sumo2* KO CD8^+^ T cells

snRNA-seq analysis was carried out for the B16(OVA) melanoma model treated with *Sumo2* KO (n = 2) or WT (n = 2) OT1 T cells using the tumor embedded Formalin-Fixed, Paraffin-Embedded (FFPE) blocks. Copy number variation analysis was conducted to identify malignant cells and other cell types were identified based on the expression of marker genes (**Fig. S9a, b, Table S1**). From the UMAP and proportion analysis (**Fig. 5A, B**), we found significantly more CD8^+^ T cells in the tumor in the mice treated with *Sumo2* KO OT1 T cells, consistent with findings using flow cytometry and IF (**Fig. 4, Fig. S9c**). Additionally, we observed an increase in dendritic cells and macrophages (**Fig. 5B, Fig. S9c**). The *Sumo2* KO CD8^+^ T cells showed enhanced expression of activation marker, (*Cd69*), co-stimulatory markers (*Cd28, Tnfrsf4*), and those reflecting effector functions (*Gzmb* and *Ifng*) compared to WT control (**Fig. 5C**). This is consistent with enhanced activation upon TCR stimulation that we observe *in vitro* (**Fig. 2**).

**Figure 5.**
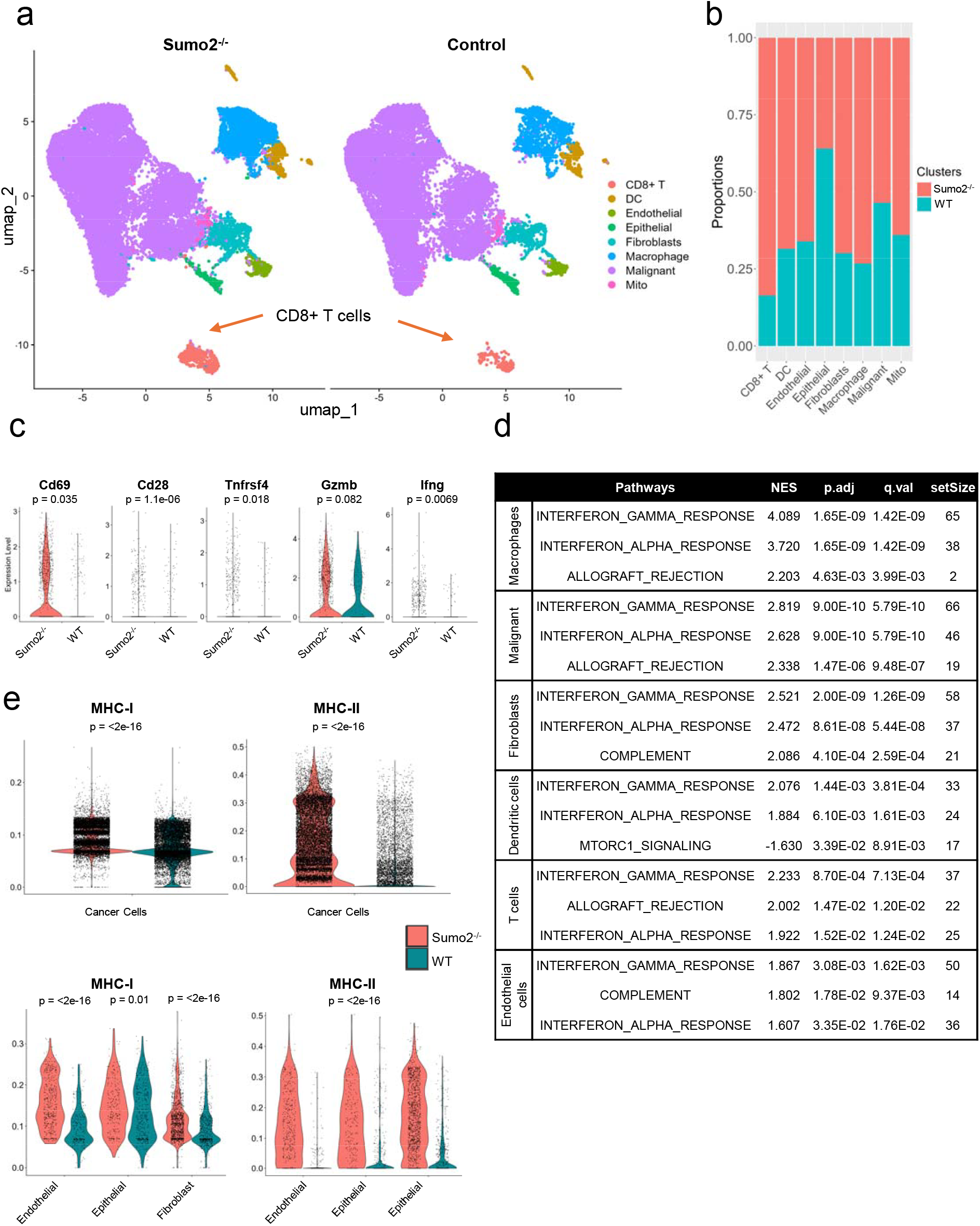
Single nuclei RNA-seq analysis reveals pathways activated *in vivo* different from those activated *in vitro*. **(A)** UMAP projections of B16(OVA) melanoma tumor samples from *Rag1*^−/−^ mice treated with *Sumo2*^−/−^ (n=2) or WT (n=2) OT-I CD8^+^ T cells. Malignant cells were determined based on abnormal copy number variation analysis from InferCNV (Figure S9A). Immune and non-immune cells were then clustered based on known cell type (Figure S9B). **(B)** Proportion analysis of the compositions of each cluster/cell type based on either *Sumo2*^−/−^ (n=2) or WT (n=2). Proportion analysis from individual replicates are shown in Figure S9C. **(C)** Violin plot of expression of activation (Cd69), co-stimulatory (Cd28) and effector function (Gzmb, Ifng) genes in CD8^+^ T cell clusters from either *Sumo2*^−/−^ or WT. Statistical significance between the two conditions was assessed using a Mann-Whitney U test. **(D)** Gene set enrichment analysis (GSEA) of hallmark gene sets conducted with clusterProfiler of differentially expressed genes in all cell types between *Sumo2*^−/−^ or WT conditions. **(E)** Violin plot of the UCell composite score of MHC-I or MHC-II (Table S1) genes expressed in malignant cells (top) or non-immune cells (bottom) in either *Sumo2*^−/−^ or WT. Statistical significance between the two conditions was assessed using a Mann-Whitney U test. **(F)** Violin plot of the UCell composite score of *Cxcl9* or *Cxcl10* genes expressed in various cell types in either *Sumo2*^−/−^ or WT. Statistical significance between the two conditions was assessed using a Mann-Whitney U test.

We conducted GSEA analysis on all major cell types present in our sample. We observed IFNγ responses in all cell types (**Fig. 5D**). We also observed IFN-I responses enriched in all cell types in tumors of mice adoptively transferred with *Sumo2* KO OT1 cells in comparison to mice adoptively transferred with control WT OT1 cells. This was unexpected, because other than CD8^+^ T cells, all immune and non-immune cells were from *Rag1*^-/-^ mice with identical genetic backgrounds. The activation of IFN-I responsive genes in *Sumo2* KO CD8^+^ T cells contrasted with these same cells when stimulated *in vitro* that had reduced expression of IFN-I responsive genes (**Fig. S10**).

MHC-I and MHC-II gene expressions on tumor cells are essential for T cell recognition in adaptive anti-tumor immune responses. Both MHC-I and MHC-II expressions increased in both the malignant cells and non-immune stromal cells including endothelial, epithelial and fibroblasts in the TME of mice adoptively transferred with *Sumo2* KO CD8^+^ T cells **(Fig. 5E)**. The expression of MHC-II is particularly pronounced in the *Sumo2* KO group **(Fig. 5E)**. The increase in both MHC-I and MHC-II are consistent with an activation of IFN-I and IFNγ signaling in these cells, as both MHC-I and MHC-II are IFN-induced genes (*15, 16*).

Chemokines CXCL9 and CXCL10 are induced by IFN-I and IFNγ and attract CTL into the TME. Consistent with the activation of IFN-I and IFNγ signaling, the expression of both genes are significantly higher in various cell types in the TME of mice receiving *Sumo2* KO CD8^+^ T cells than that receiving WT control CD8^+^ T cells **(Fig. 5F)**. This finding suggests that increased infiltration of *Sumo2* KO CD8^+^ T cells into the TME is induced by the chemokines due to activation of IFN-I and IFNγ, in addition to potentially enhanced proliferation.

### Sumo2 KO in CD8^+^ T cells dramatically alter cell-cell interactions in the TME

Cell-cell interactions in the TME were analyzed using Cellchat (*17*), which quantifies cell-cell communications based on ligand-receptor expression and gene expression patterns expected from the ligand-receptor interactions. The pathways active exclusively in mice that received WT control CD8^+^ T cells or received *Sumo2* KO CD8^+^ T cells are shown in **Fig. 6A and Fig. S11**. The distinct pathways demonstrate how modulating CD8^+^ T cells alone can profoundly impact cell-cell interactions in the TME.

**Figure 6.**
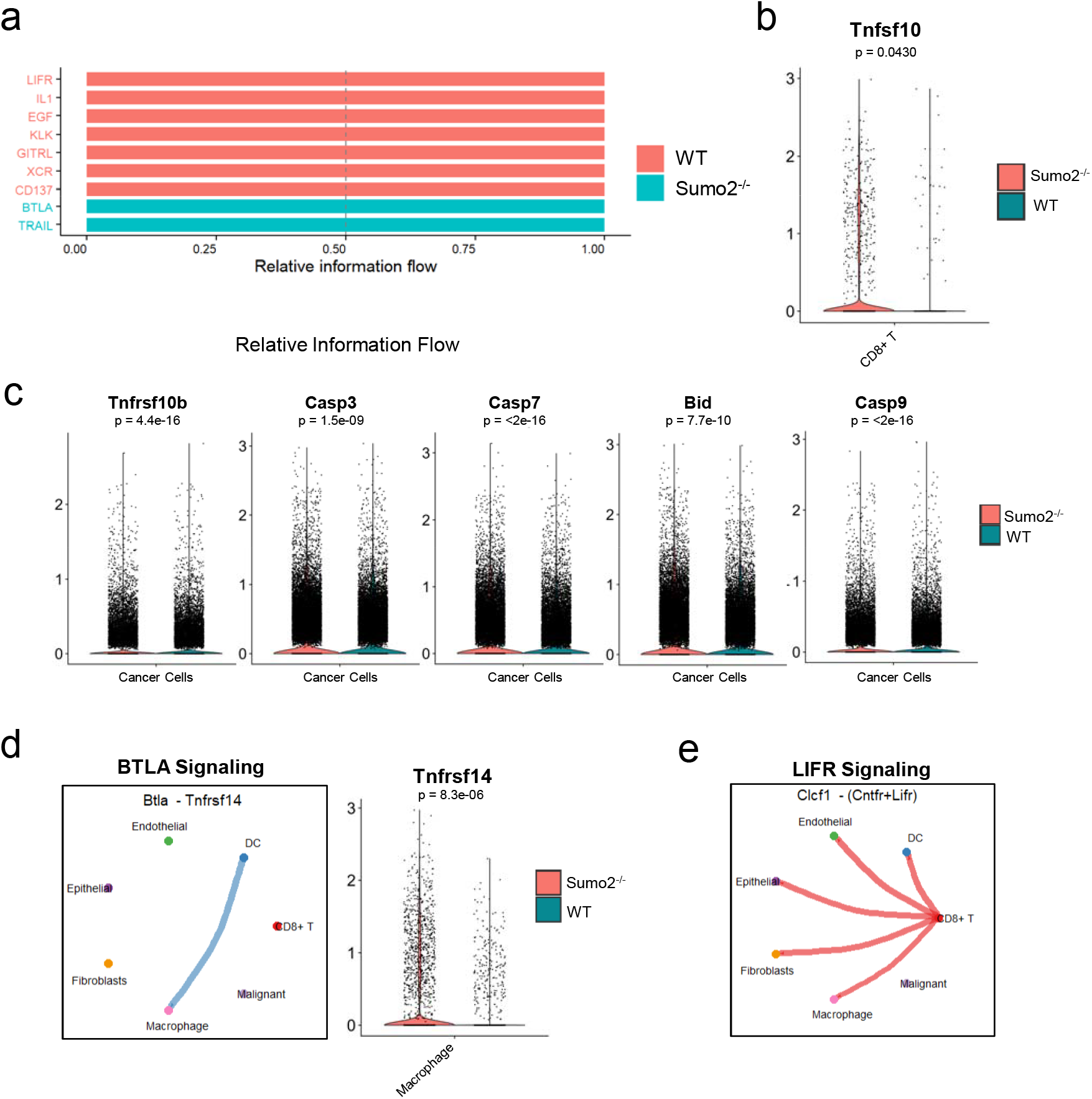
Pathways activated in tumor cells and other cells by *Sumo2*^−/−^ T cells in the tumor microenvironment. **(A)** Bar chart comparing differential pathways enriched in *Sumo2*^−/−^ or WT conditions using CellChat. Top ranking pathways from all enriched pathways (**Figure S11**) that are exclusively enriched in either condition shown. **(B)** Violin plot of expression of Tnfsf10 (TRAIL) in CD8^+^ T cells in either *Sumo2*^−/−^ or WT conditions. Statistical significance between the two conditions was assessed using a Mann-Whitney U test. **(C)**. Violin plots of expression of downstream TRAIL signaling in cancer cell populations comparing between *Sumo2*^−/−^ or WT conditions. Statistical significance between the two conditions was assessed using a Mann-Whitney U test. **(D)** Circle plot (left) showing the interacting cells, either receiving or sending, contributing to the BTLA signaling enrichment in the *Sumo2*^−/−^ condition. Violin plot (right) showing expression of Tnfrsf14 (HVEM receptor for Btla) in macrophages between either *Sumo2*^−/−^ or WT conditions. Statistical significance between the two conditions was assessed using a Mann-Whitney U test. **(E)** Circle plot showing interacting cells that contribute to the LIFR signaling pathway enrichment in the WT condition. **(F)** Schematic model of how enhanced activation of *SUMO2*^−/−^ T cells enhances tumor cell killing and alter the TME to induce increased infiltration of *SUMO2*^−/−^ T cells.

Further analysis of distinct pathways in the two groups shows that in mice receiving *Sumo2* KO CD8^+^ T cells, the TRAIL (Tnfsf10) pathway is uniquely activated. The ligand, Tnfsf10, is expressed by CD8^+^ T cells while the receptor, Tnfrsf10b, is expressed on malignant cells (**Fig. 6B**). While the expression of Tnfrsf10b was not higher in the Sumo2 KO CD8+ T cell treated condition, we observe higher levels of genes downstream of the TRAIL signaling pathway including Caspase 3, 8 and 9, and BH3 interacting domain death agonist (Bid) **(Fig. 6C)**. All of these genes are inducers of apoptosis (*18, 19*). This finding provides an understanding of improved tumor control by *Sumo2* KO CD8^+^ T cells. The *Sumo2* KO OT1 group also exhibited distinct Btla pathway activity (**Fig. 6A**). This difference stemmed from the increased expression of Tnfrsf14 (the receptor for Btla) on macrophages within the tumor microenvironment (TME) of the *Sumo2* KO group (**Fig. 6D**). Notably, the macrophages in the TME of both the *Sumo2* KO and control WT OT1 recipient mice are derived from *Rag1*^-/-^ mice and are thus genetically identical, highlighting that the adoptively transferred T cells induce this expression change. Btla has been previously identified as a lymphocyte inhibitory receptor similar to CTLA-4 and PD-1 (*20*), suggesting that this pathway may limit the anti-tumor activity of *Sumo2* KO CTL.

The pathways distinctively active in the control WT group generally are involved in tumor progression. The LIF (Leukemia Inhibitory Factor) pathway activity is predominantly due to the interaction of CD8^+^ T cells with nearly all cell types in the TME (**Fig. 6E**). This pathway has been found to stimulate cancer cell migration and is associated with unfavorable prognosis in melanoma and other cancer types (*21*). Egfr mediated pathway contributes to tumor growth and is a well-validated target for cancer therapy. Dysregulation of Kallikrein-related peptidase (KLK) pathway is also linked to cancer progression (*22*). These tumor promoting pathways were “turned off” and instead cell death pathway in tumor cells were “turned on” by *Sumo2* KO in CD8^+^ T cells.

## Discussion

Our studies described here reveal that the dominant SUMO paralog, SUMO2, directly controls cytotoxic T cell activation and effector function, likely through controlling chromatin accessibility of critical transcription factors (TF) involved. *SUMO2/Sumo2* KO increased the accessibility of the binding sites for TF important for activation and proliferation, including BATF, JunB, ATF3, FRA1, FRA2, and AP1, particularly at distal enhancers (**Fig. 2 and 3**). The BATF motif was the most significantly enriched motif in ATAC-sequence analysis of both mouse and human CD8^+^ T cells upon *SUMO2*/*Sumo2* KO. This is noteworthy because CRISPR screens have shown that BATF is a potent and central regulator of cytotoxic T cell proliferation and effector function (*12-14*). Jun and AP-1 have also been shown to play an important role in T cell activation, effector function, and suppression of exhaustion (*23, 24*). On the other hand, ETS motifs were found to be enriched in control WT T cells (**Fig. 3b**). It was shown previously that increased expression of BATF prevented T cell terminal exhaustion through counteracting Ets1 (*12*), consistent with that BATF motif is enriched in *Sumo2/SUMO2* KO T cells while Ets1 motif is enriched in WT T cells (**Fig. 3b**). In addition, WT human CD8^+^ T cells showed enrichment of LEF1 and TCF7 (a.k.a. TCF1) motifs, both of which are associated with CD8^+^-lineage T cell development (*25*). A potential future study is to perform chromatin binding studies for all of these TFs. It is possible that more than one of the TFs are involved.

The finding that SUMO2 plays an important role in regulating chromatin accessibility is supported by previous studies. SUMOylation predominantly occur in the nucleus and a major function of this process is the regulation of gene expression through modification of transcription factors and chromatin-remodeling complexes (*26-28*). Nearly all chromatin-remodeling enzymes and complexes are targets of SUMOylation. Future studies are needed to determine whether SUMOylation of a specific chromatin remodeling enzyme/complex plays a dominating role in regulating chromatin accessibility. It is likely that SUMOylation of a group of proteins, instead of one, is involved; the concept of protein group SUMOylation in regulating a particular cellular activity has been previously demonstrated (*29*).

Cell-cell interaction analysis reveals activation of TRAIL pathway through interactions between *Sumo2* KO CD8^+^ T cells and tumor cells that activated caspases and Bid in tumor cells (**Fig. 6**). TRAIL was discovered as an inducer of apoptosis that selectively targets cancer cells while leaving normal cells unharmed, because TRAIL receptors, DR4 (TRAIL-R1) and DR5 (TRAIL-R2), are often upregulated on cancer cells, while generally expressed at low levels or tightly regulated in normal cells (*30*). Therefore, delivering TRAIL and reducing its rate of elimination are being explored as a cancer therapeutic strategy (*30*). Our data has shown that *Sumo2* KO CD8^+^ T cells in the TME express higher levels of TRAIL that WT CD8^+^ T cells. In addition, expression of caspases and Bid in cancer cells was increased in cancer cells in the TME of mice received *Sumo2* KO CD8^+^ T cells indicating activation of apoptosis pathway in cancer cells (**Fig. 6**). It is of note that increased TRAIL expression by *Sumo2* KO CD8^+^ T cells only occurred *in vivo* and not *in vitro* (**Fig. S10**), consistent with differences in IFN-I and IFNγ pathway activation *in vivo* and not *in vitro*.

Our data reveals a mechanism of activating IFN-γ and IFN-I responsive genes in the TME that is independent of the known mechanism – inducing of IFN-I expression in myeloid cells by suppressing SUMOylation (**Fig. 6f**). *Sumo2* KO in CD8^+^ T cells suppressed IFN-I responsive genes in the absence of TME (**Fig. S10**). However, when *Sumo2* KO CD8^+^ T cells were adoptively transferred into tumor-bearing *Rag1*^-/-^ mice, IFN-I and IFN-γ responsive genes are activated not only in the CD8^+^ T cells, but also in all other cell types in the TME (**Fig. 5**). The enhanced activation and effector functions of the *Sumo2* KO CD8^+^ T cells could induce the expression of IFN-γ when binding to antigen-presenting tumor cells. In addition, the enhanced apoptosis of the tumor cells through TRAIL and antigen-specific killing of tumor cells could result in releasing factors to the TME to activate IFN-I (**Fig. 6f**). The enhanced IFN-I and IFN-γ response in the TME in the *Sumo2* KO group is consistent with increased MHC-I and MHC-II on both tumor cells and myeloid cells (**Fig. 5e, Fig. 6f**). Such increase in MHC-I and MHC-II are expected to enhance T cell mediated tumor killing. Activation of IFN-γ and IFN-I would induce the expression of chemokine CXCL9 and CXCL10 to enhance T cell infiltration into the TME that was observed on both the CAR-T and OT1 model (**Fig. 4, Fig. 5f, Fig. 6f**).

Our findings reveal a novel mechanism regarding how suppressing SUMOylation in CTL leads to increased CTL tumor infiltration and activation, and activates IFN-g and IFN-I responsive genes including MHC-I and MHC-II on tumor cells in the TME, phenotypes observed in the monotherapy clinical trial of TAK-981 (*9*). Similar observations have been observed in preclinical models of cancers (*15, 31*). It was shown that inhibiting SUMOylation in T cells derived from patients with chronic lymphocytic leukemia (CLL) induced T-cell activation in response to TCR stimulation and enhances secretion of IFN-γ by CD4^+^ and CD8^+^ T cells(*32*). Although these observations are similar to *SUMO2* KO shown in our studies, the previous studies suggested that the effect is mediated by IFN-I paracrine signaling, different from what we found through genome-wide analysis.

Our findings suggest that deletion of the *SUMO2* genes in T cells could be a strategy for enhancing the efficacy of adoptive T cell therapy for solid tumors, whether in the CAR-T or TCR-T format. In fact, a recent study has shown that combination of TAK-981 and 5-Aza-2’ deoxycytidine synergizes with TCR therapy in eradication of acute myeloid leukemia and multiple myeloma in animal models (*33*). This is through strong expansion of T cells and optimization of T-cell tumor cell interaction, a phenotype similar to that *SUMO2* KO in T cells as we show here, supporting our findings. snRNA-seq revealed Btla pathway as a potential inhibitory mechanism. Agents that block the Btla pathway are in early phase clinical development (*34*). Future studies are needed to determine whether blocking Btla pathway would enhance the efficacy of *SUMO2* KO T cells. Our findings provide a mechanistic rationale for *SUMO2* KO or SUMOylation inhibition as a potential strategy to enhance the effectiveness of adoptive T cell therapy.

In conclusion, SUMOylation inhibition has been established as a new clinical stage strategy to activate anti-tumor immunity (*9*). Our studies provide important new mechanistic insights into the role of SUMOylation in directly regulating CTL activation, proliferation, tumor infiltration and modulating the TME that has significant implications for developing combination strategies for effective cancer therapy.

## Materials and Methods

### Mouse Generation

To generate Mouse *Sumo2* loxP (SUMO2 FL/FL), a CRISPR/Cas9-based EGE method was used. Briefly, after analyzing the gene structure (Gene ID: 170930) and exon sizes, 5’ UTR, exon 1, and exon 2 were conditionally removed, with their deletion resulting in a null protein. This design enables conditional knockout of these regions via the Cre-loxP system. The 5’ loxP site was inserted approximately 2 kb upstream of the *Sumo2* promoter region, while the 3’ loxP site was placed in intron 2–3. This intron’s large size ensured that loxP insertion would not interfere with mRNA splicing. Both loxP sites were positioned in non-conserved regions to minimize disruption of *Sumo2* expression.

Cas9/sgRNA plasmids were constructed and validated by DNA sequencing. Plasmid efficacy was assessed using UCATM (Universal CRISPR Activity Assay), an sgRNA activity detection system (Biocytogen, China), to select constructs with higher Cas9/sgRNA activity. The selected construct was microinjected into C57BL/6N zygotes, and resulting mice were confirmed by PCR genotyping and sequencing.

B6.Cg-Tg(Lck-icre)3779Nik/J, C57BL/6-Tg(TcraTcrb)1100Mjb/J, and B6.129S7-*Rag1*^*tm1Mom*^/J mice were purchased from Jackson Laboratory. *Sumo2* FL/FL mice were bred with B6.Cg-Tg(Lck-icre)3779Nik/J (referred to as LCKcre+) to generate mice with conditional SUMO2 knockout (KO) in T cells (SUMO2 FL/FL LCKcre+). Mice with the SUMO2 ^FL/FL^ LCKcre+ genotype were then bred with C57BL/6-Tg(TcraTcrb)1100Mjb/J mice to generate mice with SUMO2KO CD8+ OT1 T cells.

### Cell lines

B16F0hCD19 cells (modified to express human CD19) and B16(OVA) cells (modified to express ovalbumin) were gifted by the Anjana Rao lab at the La Jolla Institute for Immunology, La Jolla, USA. The Plat-E retroviral packaging cell line was purchased from ATCC. These cells were cultured in DMEM supplemented with 10% (v/v) FBS, 1% (v/v) L-glutamine, and 1% (v/v) penicillin/streptomycin, and maintained at 37□°C in a 5% CO_2_ incubator. Plat-E cells were cultured with an additional supplement of 1% β-mercaptoethanol.

Jurkat cells (ATCC) were recovered from liquid nitrogen cell storage and cultured in RPMI medium supplemented with 10% (v/v) FBS, 1% (v/v) L-glutamine, and 1% (v/v) penicillin/streptomycin.

### Primary T cell isolation and culturing

Total mouse T cells or CD8^+^□T cells were isolated from the spleens of littermate control mice (SUMO2^FL/+^, SUMO2^FL/FL^ or LCKcre+) and SUMO2 KO mice using the Dynabeads™ Untouched™ Mouse T Cells Kit or Dynabeads™ Untouched™ Mouse CD8 T Cells Kit. T cells were isolated from mice aged 8–12 weeks.

A healthy human blood sample was purchased from the San Diego Blood Bank. The blood was mixed at a 1:1 ratio with PBS (2% FBS) and layered onto Lymphoprep™ solution before centrifugation at 800□g for 30 minutes. The PBMC layer was collected and washed with PBS at 1000□g for 10 minutes. Human CD8^+^ T cells were isolated using the Dynabeads™ Untouched™ Human CD8 T Cells Kit. All isolated primary T cells were cultured in RPMI medium supplemented with 10% (v/v) FBS, 1% (v/v) L-glutamine, 1% (v/v) penicillin/streptomycin and 1% β-mercaptoethanol.

### Western blot

Cells were mixed with NuPAGE LDS sample buffer (1X) and sonicated for cell lysis. The lysates were heated at 95□°C for 15 minutes. Samples were loaded into a NuPAGE 4–12% Bis-Tris gel and run at 120 V for 2 hours. The gel was blotted and transferred onto a nitrocellulose membrane at 120 V for 90 minutes at 4□°C. The membrane was washed with TBST (1X), blocked with Intercept® (TBS) blocking buffer (LI-COR Biosciences) for 1 hour, and incubated overnight at 4□°C with the primary antibody. After washing the membrane with TBST, fluorescence labelled secondary antibodies were added for 1 hour, followed by additional washing steps. Images were acquired using the LICOR ODYSSEY® CLx instrument.

### Real-time RT–qPCR

Total RNA was isolated from cells using the RNeasy Mini Kit (Qiagen) according to the manufacturer’s protocol. Total RNA concentration was determined using a NanoDrop (Invitrogen) by measuring absorbance at 260/280 nm, and 1□μg of RNA was used for reverse transcription with the PrimeScript™ 1st Strand cDNA Synthesis Kit (Takara Bio). For qPCR, 100□ng of cDNA from each sample was amplified with gene-specific primers using Power SYBR Green PCR Master Mix (Applied Biosystems) and QuantStudio□5 Real-Time PCR Systems (Applied Biosystems). Results are presented as 2^−ΔΔCt^ using reference genes.

### Flow cytometry analysis

All flow cytometry antibody staining was carried out after staining with a viability dye together with blocking antibodies. To detect intracellular markers, cells were fixed and permeabilized with the Foxp3/Transcription Factor Staining Buffer Set (eBioscience) according to the staining protocol. All samples were analyzed using Agilent NovoCyte Advanteon Flow Cytometer Systems.

### Mouse CAR T and OT1 T cell preparation for adoptive transfer

All primary T cells were cultured in RPMI medium supplemented with 10% (v/v) FBS, 1% (v/v) L-glutamine, 1% β-mercaptoethanol, and 1% (v/v) penicillin/streptomycin in 10% CO_2_ incubators. For CAR-T generation, freshly isolated CD8^+^ T cells from *Sumo2* KO or wild-type (WT) control mice were activated with 1□μg/mL anti-CD3, 1□μg/mL anti-CD28, and 100□U/mL IL-2 for 24–30 hours. Activated *Sumo2* KO and WT CD8^+^□T cells were transduced with retrovirus containing the CAR construct in 24-well plates using 1 mL of virus and 8 μg/mL of polybrene per well. Cells were spun at 37□°C for 90 minutes by centrifugation at 2000 g. Immediately after, the virus was removed and the cells were resuspended in media containing 1□μg/mL anti-CD3, 1□μg/mL anti-CD28, and 100□U/mL IL-2. A second transduction was performed one day after the first, followed by replacement of the media with 100□U/mL IL-2. CAR CD8^+^□T cells were cultured until five days post-activation. CAR T cell percentages were determined by flow cytometry, and cells were counted using a cell counter (Countess 3, Life Technologies).

Similarly, OT-I CD8^+^ T cells from *Sumo2* KO mice were isolated, activated, and cultured for seven days post-activation. During activation and culturing OT1 CD8^+^ T cells, H-2Kb OVA Tetramer-SIINFEKL-PE (5ug/ml) were added to the cell media. Cells were washed with PBS, and five million CAR T cells or OT1 T cells in 100□μL PBS were adoptively transferred to each mouse by retro-orbital intravenous injection.

### In vitro cytotoxic assay

The cytotoxicity assay was carried out using the xCELLigence Real-Time Cell Analyzer (Agilent). B16F0-hCD19 cells (10,000 cells in 100□μL of medium per well) were seeded onto an E-Plate and allowed to adhere to the electrode surface inside a 5% CO_2_ incubator. Data recording started immediately, with measurements taken at 30-minute intervals throughout the experiment. After approximately 24 hours, CAR T cells were added at various effector-to-target (E:T) ratios in a volume of 100□μL, and data acquisition continued for another 24 hours.

### B16F0-hCD19 and B16(OVA) tumor models

To analyze CAR T cells and conduct survival experiments, 8–10-week-old *Rag1*^-/-_^mice were injected subcutaneously on Day 0 with 1 × 10^5^ B16-OVA-hCD19 cells or B16(OVA) cells. On Day 5, 3 million CAR-transduced CD8^+^ T cells or 5 million OT1 CD8^+^□T cells were adoptively transferred into tumor size-matched, tumor-bearing mice. Once tumors became palpable, tumor size was measured with a caliper every other day, and tumor volume was calculated as 0.5 × (length × width^2^). On Day 7 or Day 14, tumors, spleens, and blood were collected from the mice. For the survival experiments, mice were monitored until the experimental endpoint.

### CRISPR CAS9 for gene knockout in T cells

Human CD8^+_^T cells were activated for 48 hours with Dynabeads Human T-Activator CD3/CD28 beads (Gibco). The cells were then counted, washed with PBS, and resuspended in R buffer (Neon™ Transfection System 10□μL Kit, Invitrogen). Single guide RNA (Synthego, USA) and SpCas9 nuclease protein (Alt-R™ S.p. Cas9 Nuclease V3, IDT, USA) were mixed and incubated for 5 minutes, after which the cells were added to the mixture. The cell mixture was electroporated at 1600□V with three pulses of 10□ms each using the Neon™ Nucleofector (Invitrogen).

### Retrovirus preparation

The retroviral CAR vector (MSCV-myc-CAR-2A-Thy1.1) and the pCL-Eco packaging vector were obtained from the Anjana Rao laboratory at the La Jolla Institute for Immunology. Plat-E cells (3 × 10□) were seeded in 10 cm cell culture dishes and transfected using the TransIT-LT1 Transfection Reagent (Mirus Bio) according to the manufacturer’s instructions, at a ratio of 1 μg DNA to 3 μL transfection reagent. The transfection DNA mixture was prepared in 2 mL of serum-free Opti-MEM by adding the CAR vector (10 μg), the pCL-Eco packaging vector (3.4 μg), and 40 μL of TransIT-LT1 reagent. Two days post-transfection, virus-containing supernatants were collected from the Plat-E cells. Viruses were filtered with a 0.22 μm filter (Millipore) and used immediately for transduction.

### NGS data processing

Mouse CD8+ T cells were processed for sequencing 48 hours after activation and human CD8+ T cells were processed for sequencing 4 days after CRISPR knockout of SUMO2. RNA isolated from the cells and cell pellets were sent to Novogene per the company’s instructions for RNA-Seq and ATAC-Seq pipelines respectively. All reads underwent adaptor removal, read quality filtering and trimming with fastp.□

### ATAC-Seq reads processing

ATAC-Seq reads were aligned with bowtie2 to hg38 (human) or mm39 (mouse) genome depending on the sample. The output sam files from bowtie were converted, sorted based on coordinates and indexed to bam files via samtools. Bigwig files for track visualization were generated using deepTools bamCoverage. MACS3 was used for calling peaks, using the BAMPE option and calling summits for narrow peaks. A q-value cut-off of 0.05 was used and a shift and extension for size was set to correct for transposase bias (--extsize 200 –shift −100 – nomodel). Peaks from each sample were called individually and peaks that were present in replicates were kept using bedtools intersect to generate a consensus peak list. The peaks from control and Sumo2 KO conditions were then merged to create a merged peak list, which is then used for featureCounts to generate a counts table of all the reads in the peaks. The resulting counts table was then used for downstream differential peak analysis.

### ATAC-Seq Peak Analysis

The counts table from featureCounts was used for differential peak analysis with DESeq2. The differential accessible peak sets were used for downstream analysis. Motifs and annotations of the peaks were identified using findMotifsGenome.pl and annotatePeaks.pl with HOMER using the default parameters. The histone marks (H3K27Ac - ENCSR864RGX, H3K9me3 - ENCSR772HRO and H3K27me3 - ENCSR651PFT) pull-down data were obtained from publicly available data via the ENCODE project. The pull-down was done on naive CD8^+^ alpha beta activated T cells isolated from a 30 year old male donor and had been activated *in vitro* with anti-CD3 and anti-CD28 coated beads for 7 days and with 10ng/mL of IL-2 for 5 days prior to pull-down and sequencing via Mint-ChIP-seq. TOBIAS was for TF footprint analysis following the pipeline recommended by the developers. Non-redundant JASPAR2024 CORE Vertebrates motifs were used for initial binding detection. TFs highlighted by TOBIAS (-log10(p-val) above 95% quantile or top 5% differential binding score in either direction) were then subsetted and used for the final binding detection. Highlighted TFs for both conditions were then subsetted and their respective binding scores were used for the boxplot. Candidate cis-regulatory element (cCREs) annotations from the ENCODE project were obtained via the web interface SCREEN. Bedtools intersect was used to determine the annotation of each peak. The Integrative Genomics Viewer (IGV) was used for track visualizations and figures.

### RNA-Seq reads processing

Reads from bulk RNA-seq were aligned to hg38 or mm39 genome using STAR with flags that allowed for multi-mapping reads (----outFilterMultimapNmax 100-- winAnchorMultimapNmax 100). Output sam files from the STAR aligner were then converted, sorted based on coordinates and indexed into bam files via samtools. Using the bam files and gene annotation files from either hg38 or mm39, featureCounts was used to generate the count matrices for all samples.□The count matrix was used for subsequent differential gene expression analysis.

### RNA-Seq Analysis

The counts table from featureCounts was used for differential peak analysis with DESeq2. Log CPM values were obtained to make the expression dataset input file (.gct) for Gene Set Enrichment Analysis (GSEA) analysis as well as heatmap plotting. The heatmap was generated with pheatmap using the log CPM values transformed to Z-scores with hierarchical clustering. GSEA analysis was performed with either the human (H) or mouse (MH) hallmark genesets from the MSigDb Molecular Signature Database.

### Single nuclei RNA-seq (snRNA-Seq) Sample Preparation

The snRNA-seq samples were harvested from the B16(OVA) melanoma mouse model in Rag1-/-mouse treated with adoptively transferred OT1 CD8^+^ T cells that were either *Sumo2*^-/-^ (n=2) or WT (n = 2). The tumors were harvested 10 days after adoptive cell transfer and made into FFPE blocks. To ensure only tumor tissue was retained for sequencing, the tumors were hole-punched, melted, and recast into fresh FFPE blocks. Nuclei isolation was performed following the 10x Genomics Demonstrated Protocol: Sample Preparation from FFPE Tissue Sections for Chromium Fixed RNA Profiling (CG000632). Six 50 µm scrolls per sample were used. Scrolls were deparaffinized using successive xylene and ethanol washes. The rehydrated tissue pellet was subjected to dissociation using the gentleMACS Octo Dissociator with the recommended dissociation enzyme mix to release fixed nuclei. The dissociation was performed according to the manufacturer’s program. Nuclei were washed with chilled PBS, filtered and resuspended in Tissue Resuspension Buffer. Nuclei concentration and viability were determined, and the final suspension was adjusted to a target concentration of 20,000 nuclei/µL.

Libraries followed the 10x Genomics GEM-X Flex Gene Expression protocol (CG000786 RevA) for single-plexed samples with a targeted nuclei recovery of 20,000 cells/uL. Briefly, the fixed nuclei were first subjected to Probe Hybridization with a transcriptome-wide probe set, followed by Ligation and washes. The ligated product, nuclei, and GEM-X Flex hydrogel beads (containing the 10x Barcode/UMI) were partitioned into GEMs. Inside the GEMs, Reverse Transcription (RT) generated barcoded cDNA products. This barcoded cDNA was recovered, purified using SPRIselect, and then subjected to Pre-Amplification PCR. Finally, Sample Index PCR attached the Illumina P5, P7, and the sample index to form the final sequencing-ready library, followed by SPRIselect cleanup. The four single-plexed libraries were pooled and sequenced on one lane of an Illumina NovaSeq X Plus 10B flow cell. This run delivered a total of approximately 1.25 billion single reads for the pooled samples. Based on the targeted input of 20,000 nuclei per sample, the final sequencing depth achieved was an average of 312.5 million reads per sample, resulting in 15,625 reads per nuclei. Library preparation and sequencing of the libraries were conducted at the IGM Genomics Center, University of California, San Diego, La Jolla, CA.

### Single-nuclei RNAseq data analysis

Cellranger multi was used to align and filter the fastq files obtained from the GEM-X Flex sequencing. The Chromium Mouse Transcriptome Probe set v1.1.1 (mm39) and GRCm39-2024-A reference genome was used for the alignment. The resulting filtered feature matrix was then imported into R and Seurat was primarily used for subsequent analysis. Preliminary filtering based on cut-offs (200 < nFeature_RNA < 11000 & 1000 < nCount_RNA < 200000) and percent mitochondrial reads (10%) was done. Preliminary analysis including clustering was also performed to assess cell cycle gene effects and necessity for integration. Cells were normalized using SCTransform with Seurat, regressing out cell cycle genes and mitochondrial genes, and integrated for downstream analysis. Cells were clustered at a resolution of 0.4. Cell cluster markers were identified with FindAllMarkers (logfc.threshold = 0.25, min.pct = 0.25). All cluster marker genes were saved as tables (Table S1) and the cluster with predominantly mitochondrial genes were removed from downstream analysis. Obvious non-malignant clusters were identified via cell specific markers (CD8^+^ T cells, dendritic cells, fibroblasts and macrophages) and used as control references for subsequent copy number variation analysis. Copy number variation analysis was done with InferCNV for malignant cell cluster identification. Final cell types were then identified based on known markers. UCell was used for scoring and plotting the MHC-I and MHC-II expression between the two conditions in different cell types. All MHC-I and MHC-II genes detected in the dataset were compiled and used as the ‘gene signature’ for UCell scoring. Differential genes for each cluster across the two conditions were obtained using FindMarkers (min.pct = 0.25, logfc.threshold = 0.25). Statistically significant differentially expressed genes (p.adj < 0.05) and their average log2FC were used for the mouse hallmark geneset (from msigdbr) for GSEA analysis with clusterprofiler.

## Data Availability

Raw and processed sequencing data generated in this study is deposited and accessible on the NCBI Gene Expression Omnibus (GSE313614, GSE313360, GSE313361, GSE313362, GSE313363).

## Acknowledgements

We thank the Moores Cancer Center Tissue Technology Services Histology core for their assistance with FFPE embedding and IF staining. We thank Sam Parsons from 10x Genomics, Dr. Kristen Jepsen and members of the IGM Genomics Center at UC San Diego for their guidance and assistance with the sample preparation and sequencing of the snRNAseq. We thank Dr. Anjana Rao and her lab members for their guidance and assistance with the CAR-T constructs and melanoma cell lines. This publication includes data generated at the UC San Diego IGM Genomics Center utilizing an Illumina NovaSeq X Plus that was purchased with funding from a National Institutes of Health SIG grant (#S10 OD026929).

## References

1. A. Decque et al., Sumoylation coordinates the repression of inflammatory and anti-viral gene-expression programs during innate sensing. Nat Immunol 17, 140–149 (2016).

2. J. Song, L. K. Durrin, T. A. Wilkinson, T. G. Krontiris, Y. Chen, Identification of a SUMO-binding motif that recognizes SUMO-modified proteins. Proceedings of the National Academy of Sciences of the United States of America 101, 14373–14378 (2004).

3. J. Song, Z. Zhang, W. Hu, Y. Chen, Small ubiquitin-like modifier (SUMO) recognition of a SUMO binding motif: a reversal of the bound orientation. The Journal of biological chemistry 280, 40122–40129 (2005).

4. J. S. Seeler, A. Dejean, SUMO and the robustness of cancer. Nat Rev Cancer 17, 184–197 (2017).

5. L. Du et al., Role of SUMO activating enzyme in cancer stem cell maintenance and self-renewal. Nature communications 7, 12326 (2016).

6. A. Biederstadt et al., SUMO pathway inhibition targets an aggressive pancreatic cancer subtype. Gut 69, 1472–1482 (2020).

7. J. D. Kessler et al., A SUMOylation-dependent transcriptional subprogram is required for Myc-driven tumorigenesis. Science 335, 348–353 (2012).

8. E. S. Lightcap et al., A small-molecule SUMOylation inhibitor activates antitumor immune responses and potentiates immune therapies in preclinical models. Sci Transl Med 13, eaba7791 (2021).

9. D. Juric et al., A First-In-Human Study of the SUMOylation Inhibitor Subasumstat In Patients With Advanced/Metastatic Solid Tumors or Relapsed/Refractory Hematologic Malignancies. Cancer Res Commun, (2025).

10. L. Wang et al., SUMO2 is essential while SUMO3 is dispensable for mouse embryonic development. EMBO Rep 15, 878–885 (2014).

11. J. Chen et al., NR4A transcription factors limit CAR T cell function in solid tumours. Nature 567, 530–534 (2019).

12. P. Zhou et al., Single-cell CRISPR screens in vivo map T cell fate regulomes in cancer. Nature 624, 154–163 (2023).

13. F. Blaeschke et al., Modular pooled discovery of synthetic knockin sequences to program durable cell therapies. Cell 186, 4216–4234 e4233 (2023).

14. J. Wei et al., Targeting REGNASE-1 programs long-lived effector T cells for cancer therapy. Nature 576, 471–476 (2019).

15. U. M. Demel et al., Activated SUMOylation restricts MHC class I antigen presentation to confer immune evasion in cancer. J Clin Invest 132, (2022).

16. D. P. Simmons et al., Type I IFN drives a distinctive dendritic cell maturation phenotype that allows continued class II MHC synthesis and antigen processing. Journal of immunology 188, 3116–3126 (2012).

17. S. Jin et al., Inference and analysis of cell-cell communication using CellChat. Nature communications 12, 1088 (2021).

18. S. Shalini, L. Dorstyn, S. Dawar, S. Kumar, Old, new and emerging functions of caspases. Cell Death Differ 22, 526–539 (2015).

19. K. Wang, X. M. Yin, D. T. Chao, C. L. Milliman, S. J. Korsmeyer, BID: a novel BH3 domain-only death agonist. Genes Dev 10, 2859–2869 (1996).

20. N. Watanabe et al., BTLA is a lymphocyte inhibitory receptor with similarities to CTLA-4 and PD-1. Nat Immunol 4, 670–679 (2003).

21. H. Guo, Y. Cheng, M. Martinka, K. McElwee, High LIFr expression stimulates melanoma cell migration and is associated with unfavorable prognosis in melanoma. Oncotarget 6, 25484–25498 (2015).

22. T. Kryza, M. L. Silva, D. Loessner, N. Heuze-Vourc’h, J. A. Clements, The kallikrein-related peptidase family: Dysregulation and functions during cancer progression. Biochimie 122, 283–299 (2016).

23. R. C. Lynn et al., c-Jun overexpression in CAR T cells induces exhaustion resistance. Nature 576, 293–300 (2019).

24. F. Macian et al., Transcriptional mechanisms underlying lymphocyte tolerance. Cell 109, 719–731 (2002).

25. S. Xing et al., Tcf1 and Lef1 transcription factors establish CD8(+) T cell identity through intrinsic HDAC activity. Nat Immunol 17, 695–703 (2016).

26. G. Gill, Post-translational modification by the small ubiquitin-related modifier SUMO has big effects on transcription factor activity. Curr Opin Genet Dev 13, 108–113 (2003).

27. E. T. Yeh, SUMOylation and De-SUMOylation: wrestling with life’s processes. The Journal of biological chemistry 284, 8223–8227 (2009).

28. R. T. Hay, SUMO: a history of modification. Molecular cell 18, 1–12 (2005).

29. I. Psakhye, S. Jentsch, Protein group modification and synergy in the SUMO pathway as exemplified in DNA repair. Cell 151, 807–820 (2012).

30. H. Walczak et al., Tumoricidal activity of tumor necrosis factor-related apoptosis-inducing ligand in vivo. Nat Med 5, 157–163 (1999).

31. S. Erdem et al., Inhibition of SUMOylation Induces Adaptive Anti-Tumor Immunity Against Pancreatic Cancer through Multiple Effects on the Tumor Microenvironment. Molecular cancer therapeutics in press, (2024).

32. V. Lam et al., T Cell-intrinsic Immunomodulatory Effects of TAK-981 (Subasumstat), a SUMO-activating Enzyme Inhibitor, in Chronic Lymphocytic Leukemia. Molecular cancer therapeutics 22, 1040–1051 (2023).

33. J. S. Kroonen et al., Targeting epigenetic regulation and post-translational modification with 5-Aza-2’ deoxycytidine and SUMO E1 inhibition augments T-cell receptor therapy. J Immunother Cancer 12, (2024).

34. Y. L. Chen et al., BTLA blockade enhances Cancer therapy by inhibiting IL-6/IL-10-induced CD19(high) B lymphocytes. J Immunother Cancer 7, 313 (2019).

