## Supplemental Figures for "SUMO2 Deletion Changes Chromatin Accessibility and Enhances Cytotoxic T Cell Activation and Tumor Infiltration"

### Slide 1
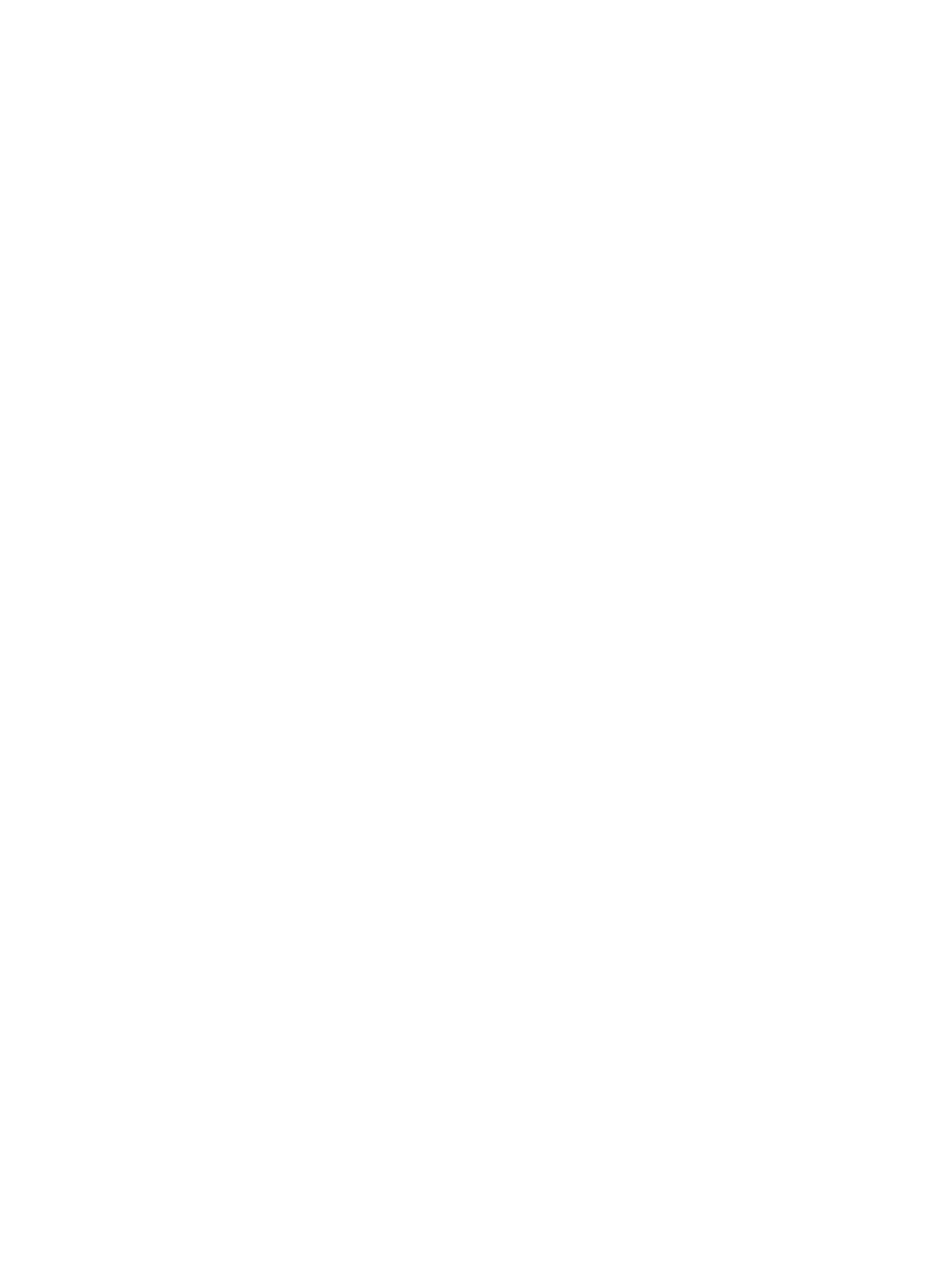

### Slide 2
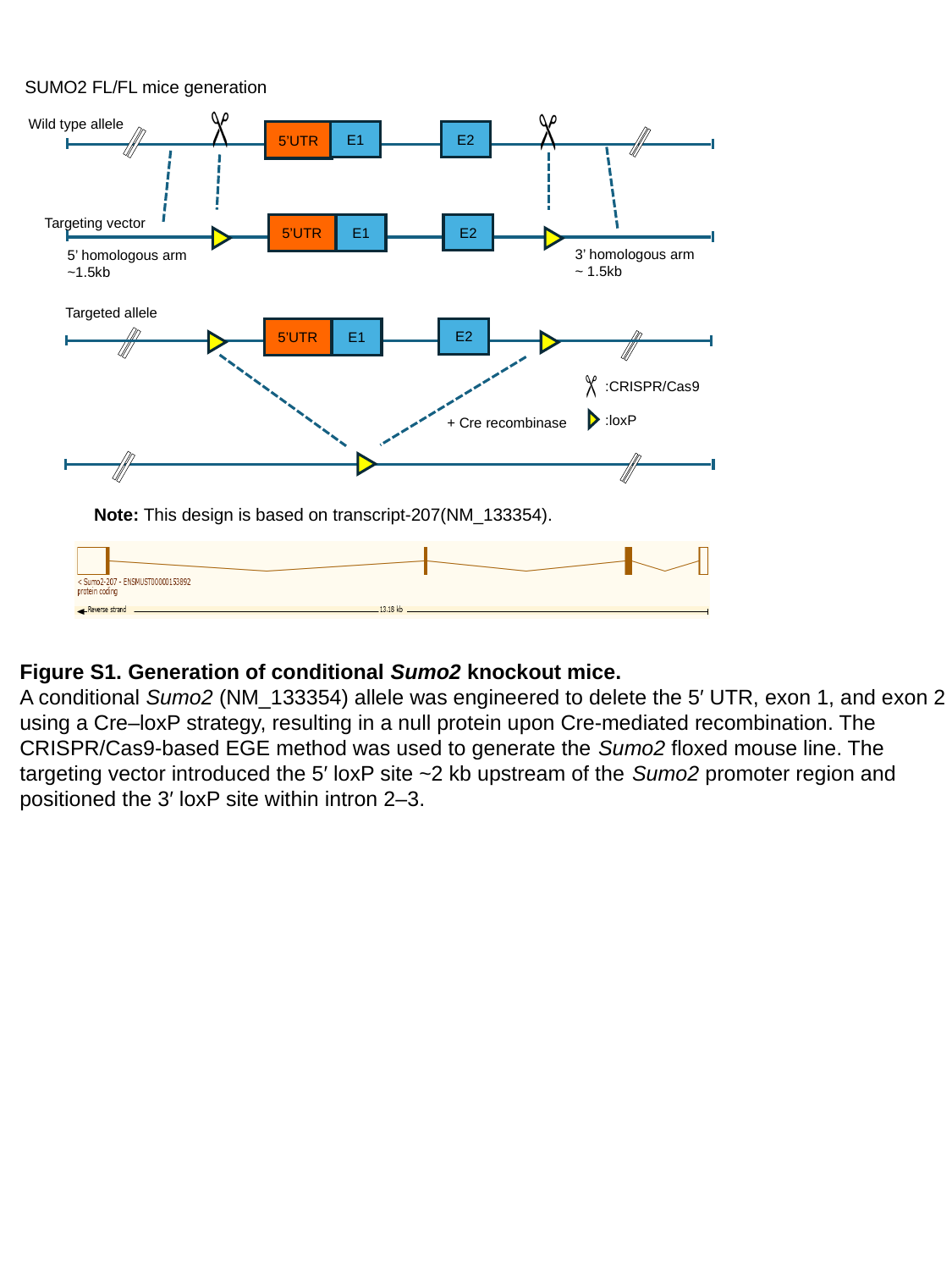

SUMO2 FL/FL mice generation
E1
E2
5’UTR
Targeting vector
E2
5’UTR
E1
3’ homologous arm
~ 1.5kb
5’ homologous arm
~1.5kb
Targeted allele
E2
5’UTR
E1
+ Cre recombinase
Wild type allele
:CRISPR/Cas9
:loxP
Note: This design is based on transcript-207(NM_133354).
Figure S1. Generation of conditional Sumo2 knockout mice.
A conditional Sumo2 (NM_133354) allele was engineered to delete the 5′ UTR, exon 1, and exon 2 using a Cre–loxP strategy, resulting in a null protein upon Cre-mediated recombination. The CRISPR/Cas9-based EGE method was used to generate the Sumo2 floxed mouse line. The targeting vector introduced the 5′ loxP site ~2 kb upstream of the Sumo2 promoter region and positioned the 3′ loxP site within intron 2–3.

### Slide 3
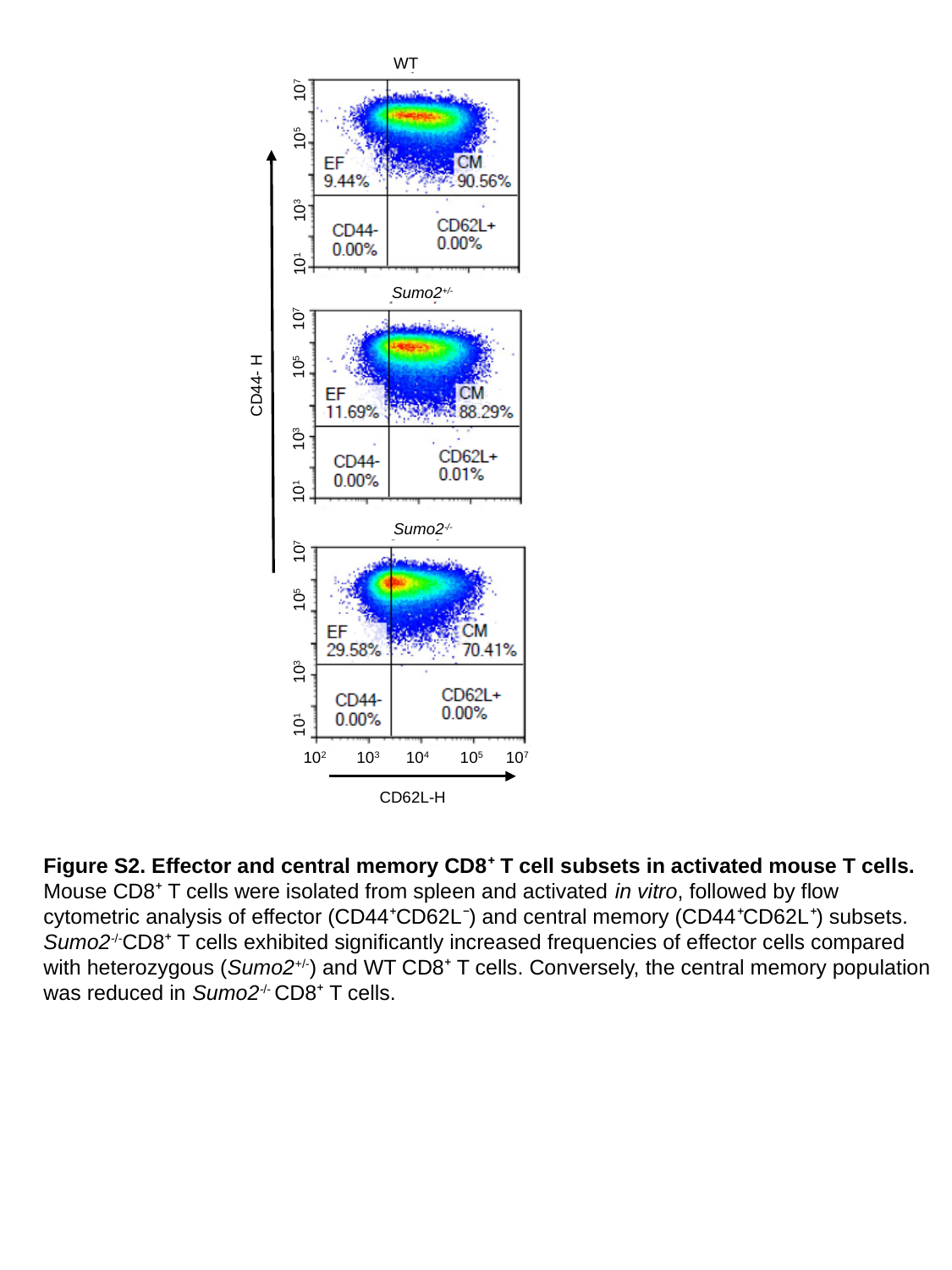

WT
 10^1^ 10^3^ 10^5^ 10^7^
CD44- H
 10^1^ 10^3^ 10^5^ 10^7^
CD62L-H
Sumo2^+/-^
Sumo2^-/-^
 10^1^ 10^3^ 10^5^ 10^7^
 10^2^ 10^3^ 10^4^ 10^5^ 10^7^
Figure S2. Effector and central memory CD8⁺ T cell subsets in activated mouse T cells. Mouse CD8⁺ T cells were isolated from spleen and activated in vitro, followed by flow cytometric analysis of effector (CD44⁺CD62L⁻) and central memory (CD44⁺CD62L⁺) subsets. Sumo2^-/-^CD8⁺ T cells exhibited significantly increased frequencies of effector cells compared with heterozygous (Sumo2^+/-^) and WT CD8⁺ T cells. Conversely, the central memory population was reduced in Sumo2^-/-^ CD8⁺ T cells.

### Slide 4
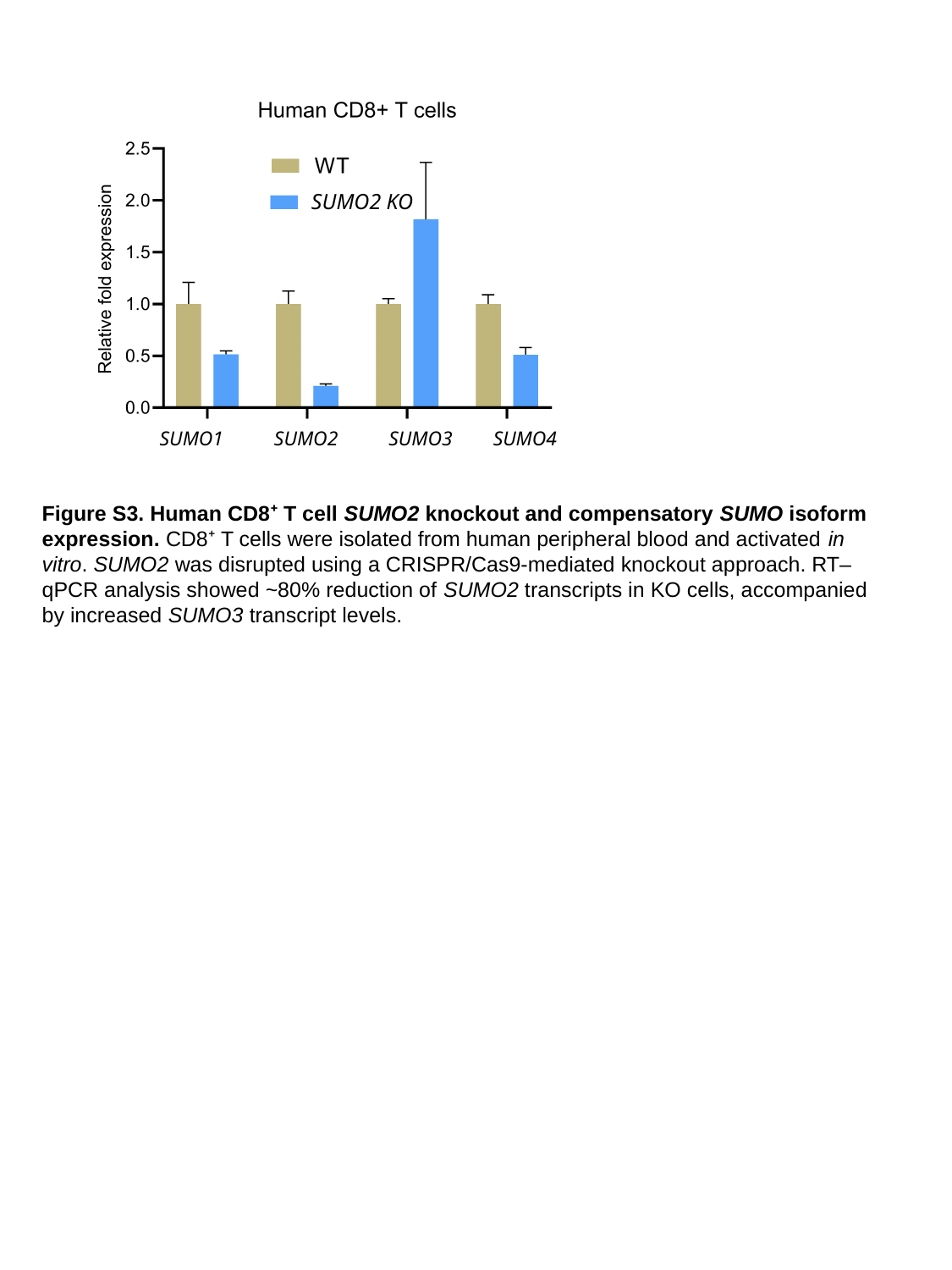

SUMO2 KO
SUMO1 SUMO2 SUMO3 SUMO4
Figure S3. Human CD8⁺ T cell SUMO2 knockout and compensatory SUMO isoform expression. CD8⁺ T cells were isolated from human peripheral blood and activated in vitro. SUMO2 was disrupted using a CRISPR/Cas9-mediated knockout approach. RT–qPCR analysis showed ~80% reduction of SUMO2 transcripts in KO cells, accompanied by increased SUMO3 transcript levels.

### Slide 5
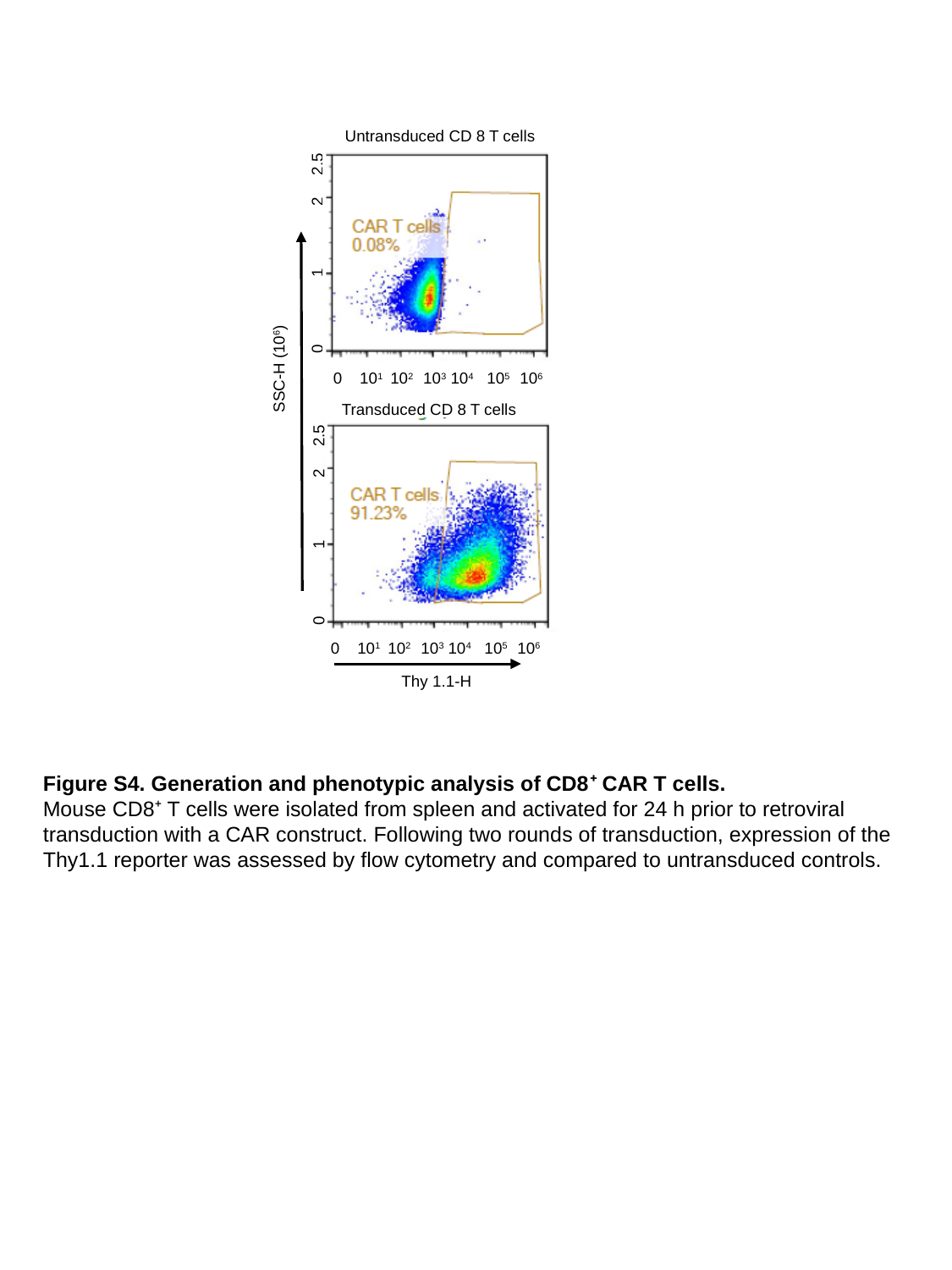

Untransduced CD 8 T cells
 0 1 2 2.5
SSC-H (10^6^)
0 10^1^ 10^2^ 10^3^ 10^4^ 10^5^ 10^6^
 0 1 2 2.5
0 10^1^ 10^2^ 10^3^ 10^4^ 10^5^ 10^6^
Thy 1.1-H
Transduced CD 8 T cells
Figure S4. Generation and phenotypic analysis of CD8⁺ CAR T cells.
Mouse CD8⁺ T cells were isolated from spleen and activated for 24 h prior to retroviral transduction with a CAR construct. Following two rounds of transduction, expression of the Thy1.1 reporter was assessed by flow cytometry and compared to untransduced controls.

### Slide 6
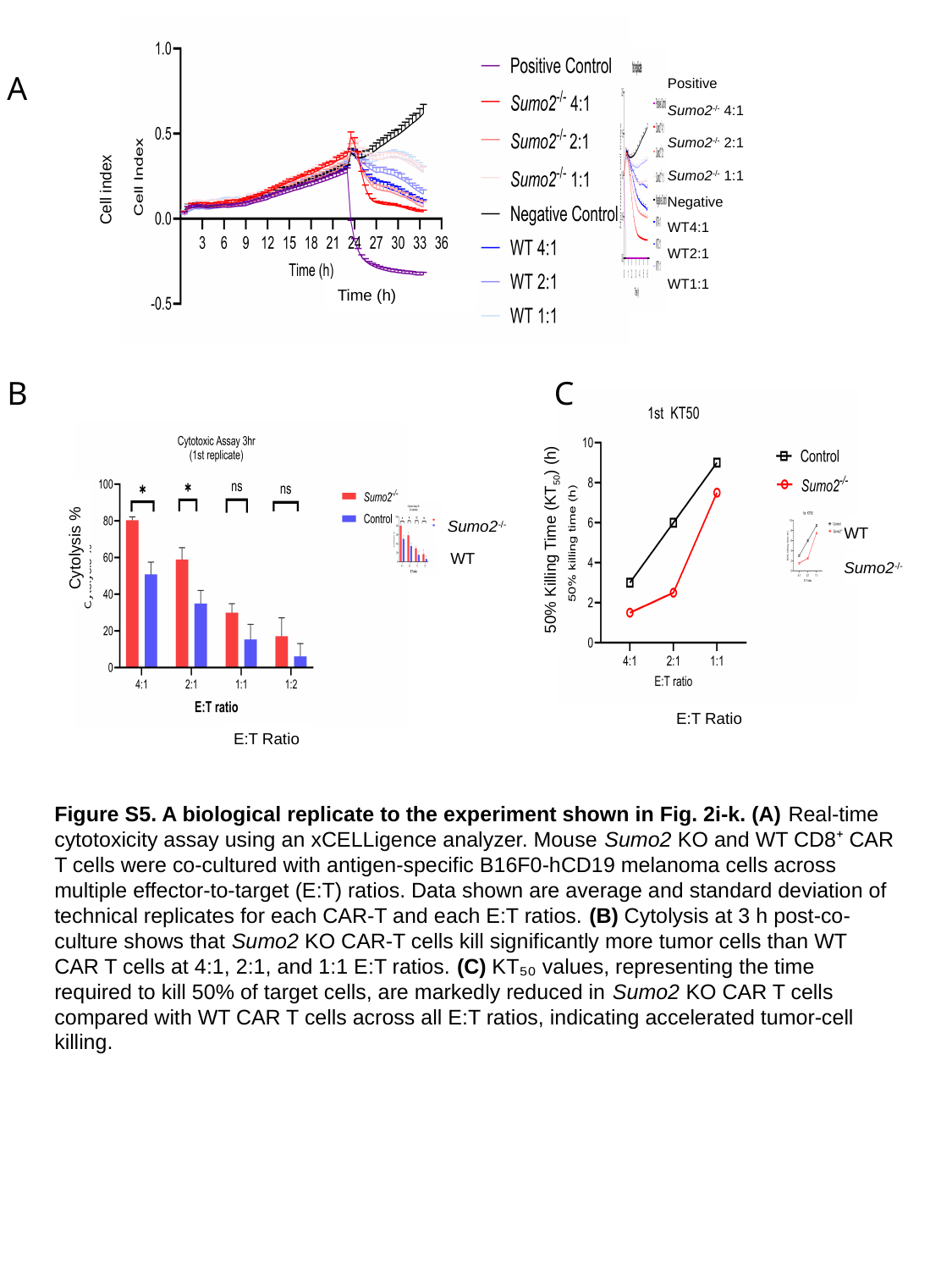

Positive
Sumo2^-/-^ 4:1
Cell index
Sumo2^-/-^ 2:1
Sumo2^-/-^ 1:1
Negative
WT4:1
WT2:1
WT1:1
Time (h)
A
B C
WT
 50% Killing Time (KT_50_) (h)
Sumo2^-/-^
E:T Ratio
Sumo2^-/-^
Cytolysis %
WT
E:T Ratio
Figure S5. A biological replicate to the experiment shown in Fig. 2i-k. (A) Real-time cytotoxicity assay using an xCELLigence analyzer. Mouse Sumo2 KO and WT CD8⁺ CAR T cells were co-cultured with antigen-specific B16F0-hCD19 melanoma cells across multiple effector-to-target (E:T) ratios. Data shown are average and standard deviation of technical replicates for each CAR-T and each E:T ratios. (B) Cytolysis at 3 h post-co-culture shows that Sumo2 KO CAR-T cells kill significantly more tumor cells than WT CAR T cells at 4:1, 2:1, and 1:1 E:T ratios. (C) KT₅₀ values, representing the time required to kill 50% of target cells, are markedly reduced in Sumo2 KO CAR T cells compared with WT CAR T cells across all E:T ratios, indicating accelerated tumor-cell killing.

### Slide 7
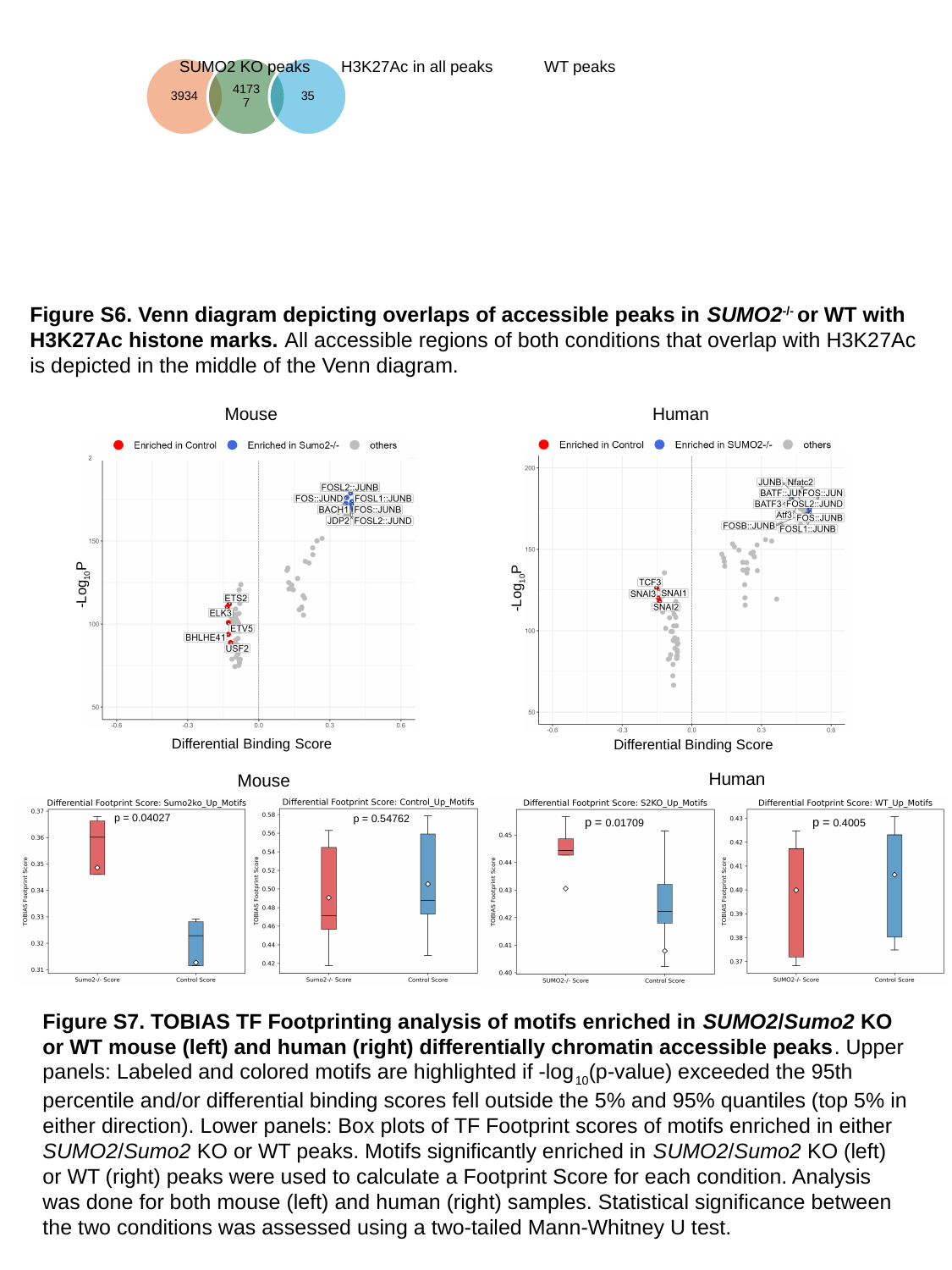

SUMO2 KO peaks
H3K27Ac in all peaks
WT peaks
127
1776
Figure S6. Venn diagram depicting overlaps of accessible peaks in SUMO2^-/-^ or WT with H3K27Ac histone marks. All accessible regions of both conditions that overlap with H3K27Ac is depicted in the middle of the Venn diagram.
Human
Mouse
Differential Binding Score
-Log_10_P
-Log_10_P
Differential Binding Score
Human
Mouse
p = 0.04027
p = 0.54762
p = 0.01709
p = 0.4005
Figure S7. TOBIAS TF Footprinting analysis of motifs enriched in SUMO2/Sumo2 KO or WT mouse (left) and human (right) differentially chromatin accessible peaks. Upper panels: Labeled and colored motifs are highlighted if -log_10_(p-value) exceeded the 95th percentile and/or differential binding scores fell outside the 5% and 95% quantiles (top 5% in either direction). Lower panels: Box plots of TF Footprint scores of motifs enriched in either SUMO2/Sumo2 KO or WT peaks. Motifs significantly enriched in SUMO2/Sumo2 KO (left) or WT (right) peaks were used to calculate a Footprint Score for each condition. Analysis was done for both mouse (left) and human (right) samples. Statistical significance between the two conditions was assessed using a two-tailed Mann-Whitney U test.

### Slide 8
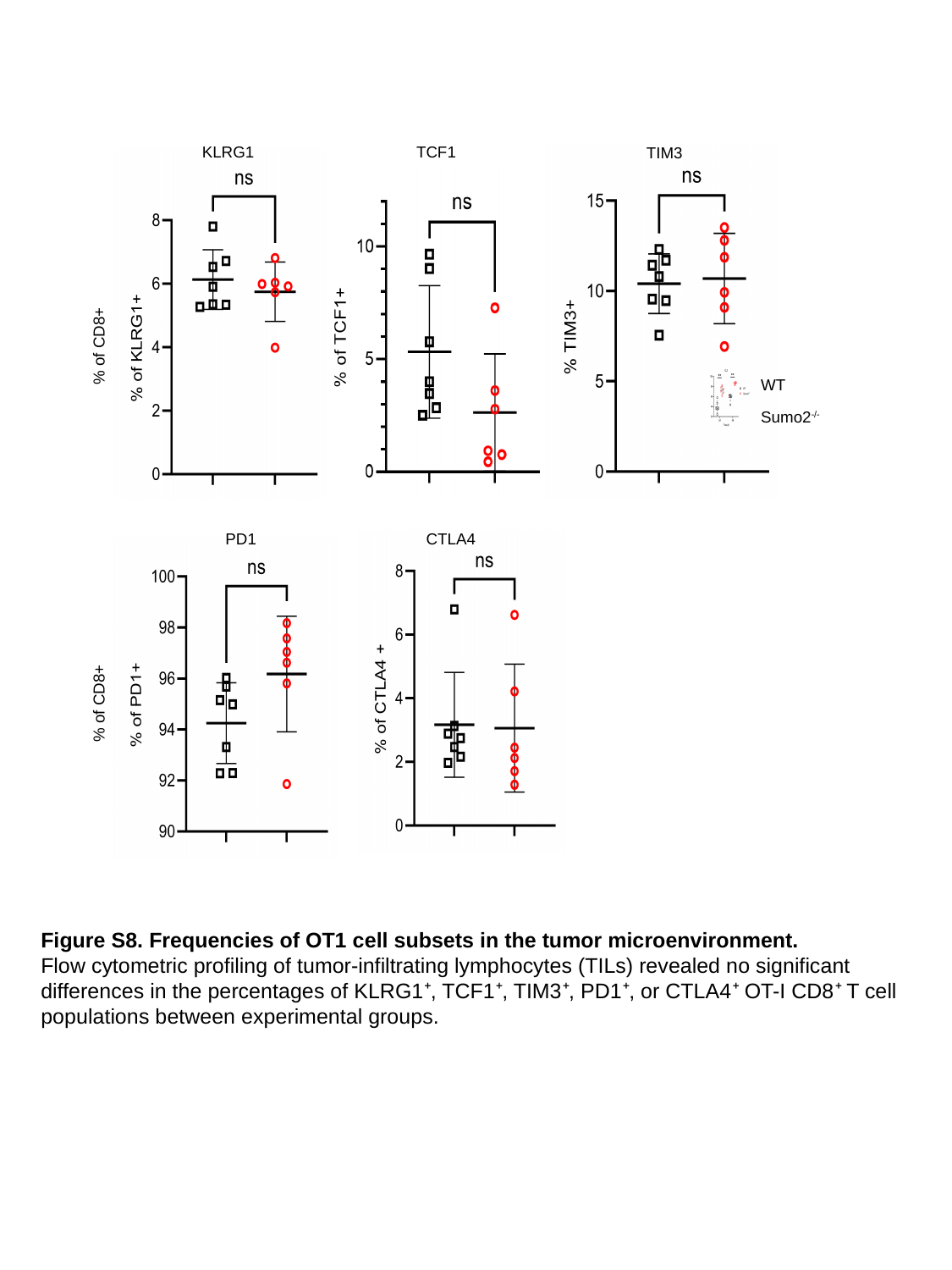

KLRG1
TCF1
TIM3
% of CD8+
WT
Sumo2^-/-^
PD1
CTLA4
% of CD8+
Figure S8. Frequencies of OT1 cell subsets in the tumor microenvironment.
Flow cytometric profiling of tumor-infiltrating lymphocytes (TILs) revealed no significant differences in the percentages of KLRG1⁺, TCF1⁺, TIM3⁺, PD1⁺, or CTLA4⁺ OT-I CD8⁺ T cell populations between experimental groups.

### Slide 9
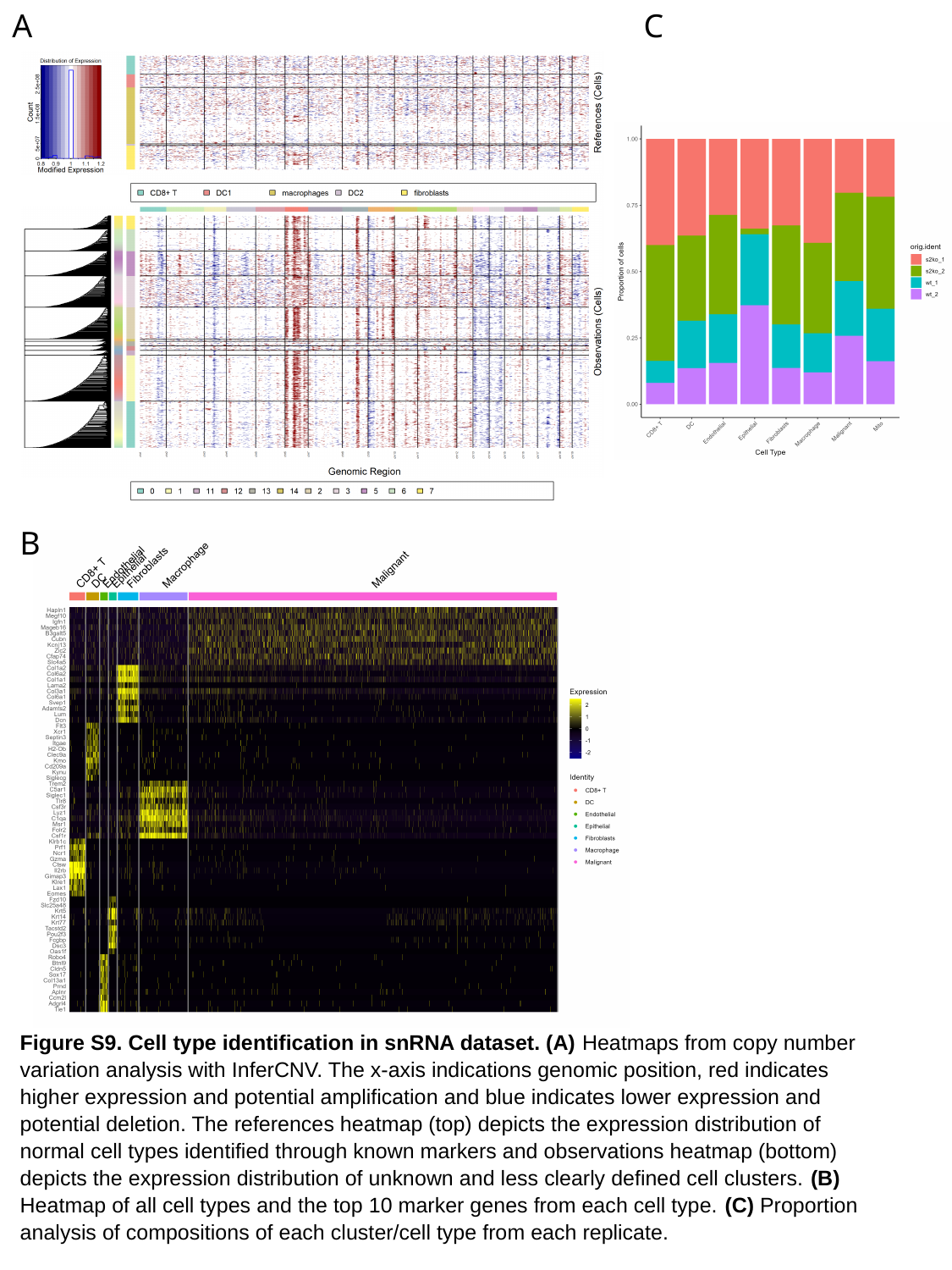

A
C
B
Figure S9. Cell type identification in snRNA dataset. (A) Heatmaps from copy number variation analysis with InferCNV. The x-axis indications genomic position, red indicates higher expression and potential amplification and blue indicates lower expression and potential deletion. The references heatmap (top) depicts the expression distribution of normal cell types identified through known markers and observations heatmap (bottom) depicts the expression distribution of unknown and less clearly defined cell clusters. (B) Heatmap of all cell types and the top 10 marker genes from each cell type. (C) Proportion analysis of compositions of each cluster/cell type from each replicate.

### Slide 10
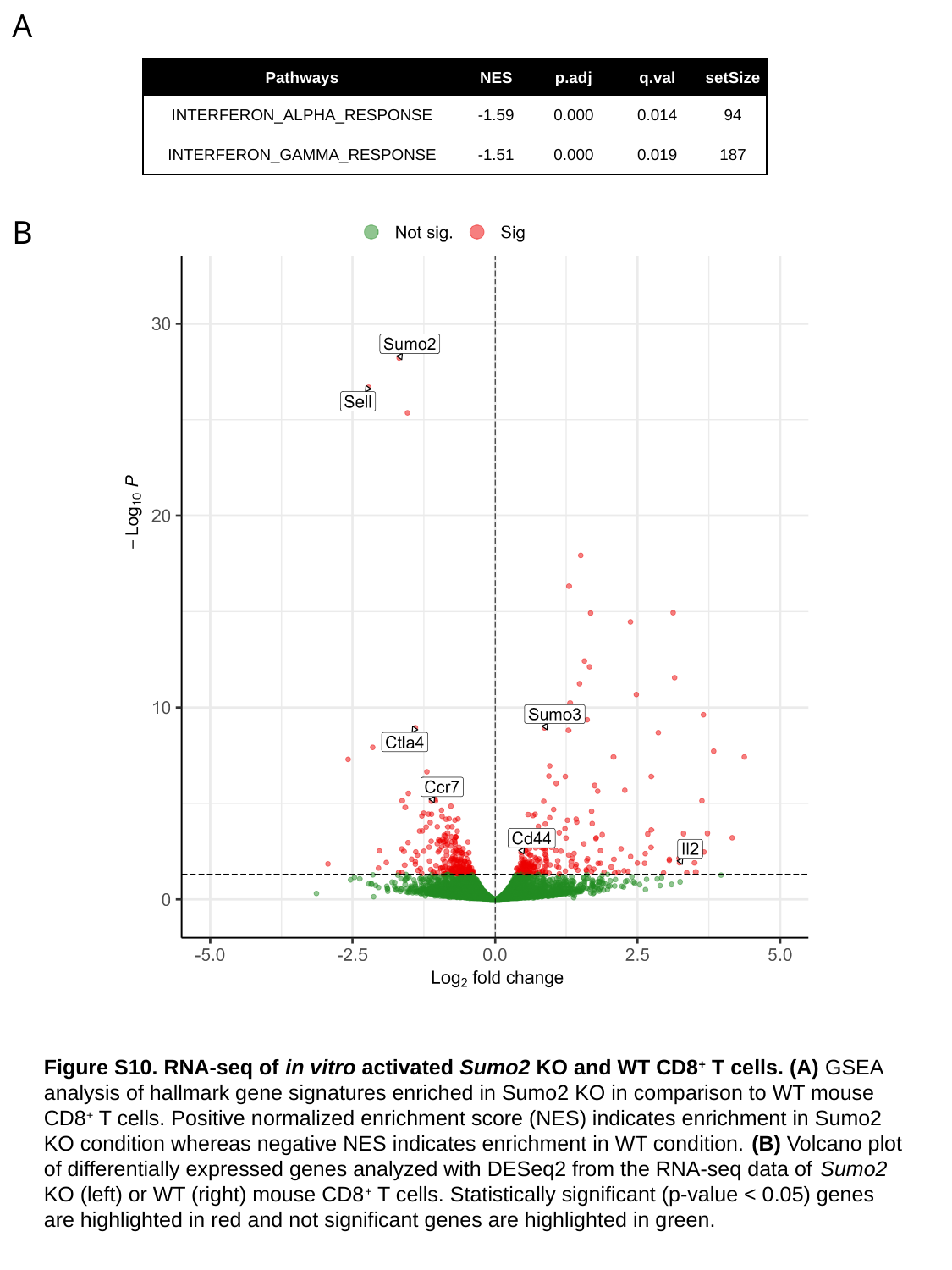

A
| Pathways | NES | p.adj | q.val | setSize |
| --- | --- | --- | --- | --- |
| INTERFERON\_ALPHA\_RESPONSE | -1.59 | 0.000 | 0.014 | 94 |
| INTERFERON\_GAMMA\_RESPONSE | -1.51 | 0.000 | 0.019 | 187 |
B
Figure S10. RNA-seq of in vitro activated Sumo2 KO and WT CD8^+^ T cells. (A) GSEA analysis of hallmark gene signatures enriched in Sumo2 KO in comparison to WT mouse CD8^+^ T cells. Positive normalized enrichment score (NES) indicates enrichment in Sumo2 KO condition whereas negative NES indicates enrichment in WT condition. (B) Volcano plot of differentially expressed genes analyzed with DESeq2 from the RNA-seq data of Sumo2 KO (left) or WT (right) mouse CD8^+^ T cells. Statistically significant (p-value < 0.05) genes are highlighted in red and not significant genes are highlighted in green.

### Slide 11
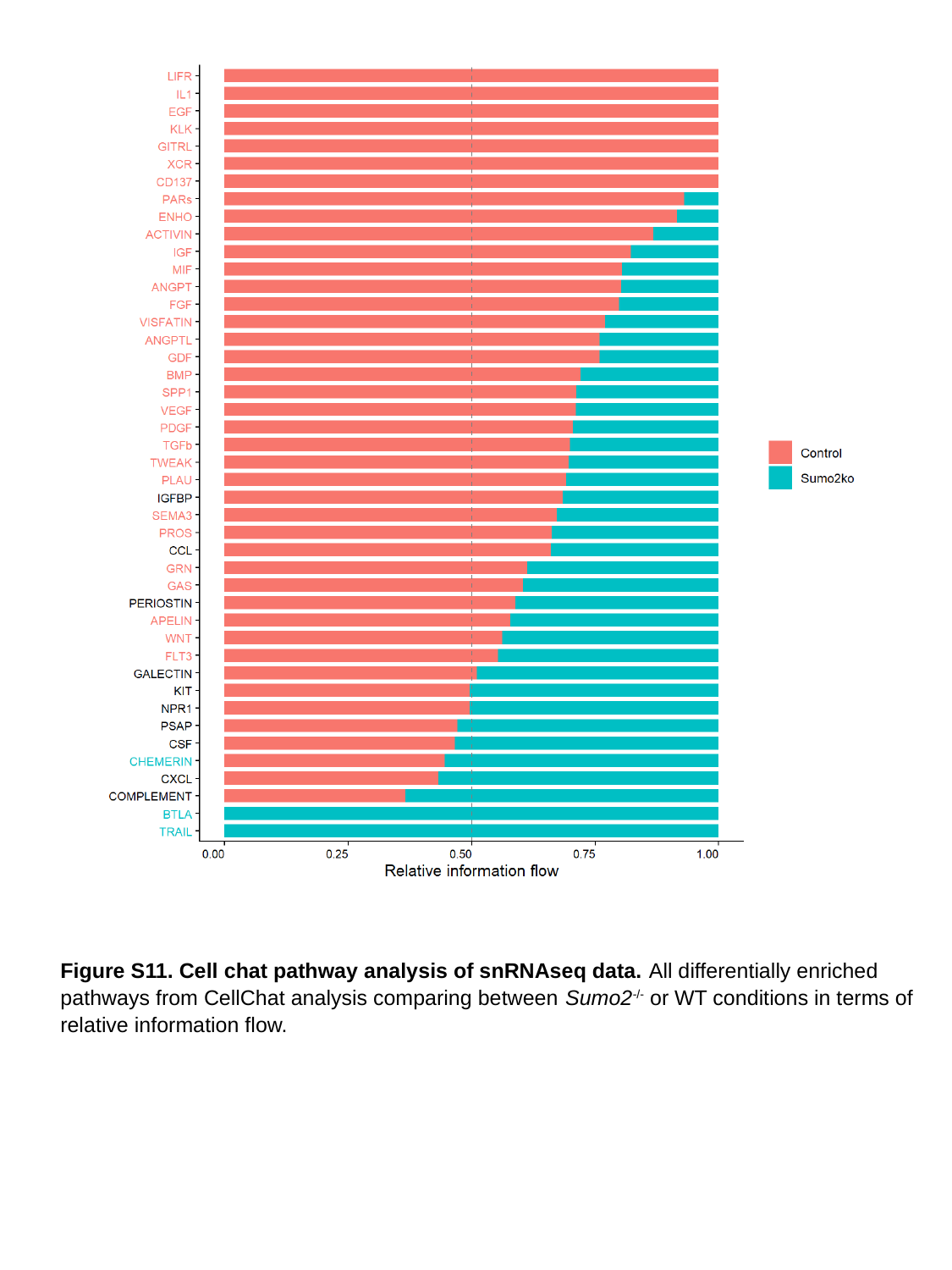

Figure S11. Cell chat pathway analysis of snRNAseq data. All differentially enriched pathways from CellChat analysis comparing between Sumo2^-/-^ or WT conditions in terms of relative information flow.
