## Supplemental Tables for "SUMO2 Deletion Changes Chromatin Accessibility and Enhances Cytotoxic T Cell Activation and Tumor Infiltration"

**Command Line Tools**

| **Package** | **Version** | **Source** |
| --- | --- | --- |
| fastp | v0.20.0 | <https://doi.org/10.1093/bioinformatics/bty560> |
| bowtie2 | v2.5.4 | <https://doi.org/10.1038/nmeth.1923> |
| MACS3 | v3.0.3 | <https://doi.org/10.1186/gb-2008-9-9-r137> |
| TOBIAS | v0.13.3 | <https://doi.org/10.1038/s41467-020-18035-1> |
| HOMER | v5.1 | <https://doi.org/10.1016/j.molcel.2010.05.004> |
| STAR | v2.7.3a | <https://doi.org/10.1093/bioinformatics/bts635> |
| SAMtools | v1.2 | <https://doi.org/10.1093/gigascience/giab008> |
| bedtools | v2.31.1 | <https://doi.org/10.1093/bioinformatics/btq033> |
| deeptools | v3.5.5 | <https://doi.org/10.1093/nar/gku365> |
| featureCounts | v2.0.0 | <https://doi.org/10.1093/bioinformatics/btt656> |
| cellranger | v9.0.1 | <https://doi.org/10.1038/ncomms14049> |

**R Packages**

| **Package** | **Version** | **Source** |
| --- | --- | --- |
| DESeq2 | v1.42.1 | <https://doi.org/10.1186/s13059-014-0550-8> |
| EnhancedVolcano | v1.20.0 | [https://github.com/kevinblighe/EnhancedVolcano](https://github.com/kevinblighe/EnhancedVolcano.) |
| pheatmap | v1.0.13 | <https://github.com/raivokolde/pheatmap> |
| msigdbr | v25.1.1 | [https://igordot.github.io/msigdbr/.](https://igordot.github.io/msigdbr/) |
| Seurat | v5.2.1 | <https://doi.org/10.1038/s41587-023-01767-y> |
| InferCNV | v1.18.1 | <https://github.com/broadinstitute/inferCNV> |
| UCell | v.2.6.2 | <https://doi.org/10.1016/j.csbj.2021.06.043> |
| ggplot2 | v.3.5.2 | [https://ggplot2.tidyverse.org](https://ggplot2.tidyverse.org/) |
| CellChat | v2.2.0 | <https://doi.org/10.1101/2023.11.05.565674> |

**Others**

| **Software/Name** | **Version** | **Source** |
| --- | --- | --- |
| JASPAR CORE | 2024 | <https://doi.org/10.1093/nar/gkad1059> |
| hg38 | GRCh38.p14 | <https://doi.org/10.1038/s41597-024-03571-y> |
| mm39 | GRCm39 | <https://doi.org/10.1038/s41597-024-03571-y> |
| ENCODE/SCREEN |  | <https://doi.org/10.1038/s41586-020-2493-4> |
| GSEA | v4.2.3 | <https://doi.org/10.1073/pnas.0506580102>, <https://doi.org/10.1038/ng1180> |
| IGV | v2.17.4 | <https://doi.org/10.1038/nbt.1754> |

**Table of antibodies**

| **Catalog No.** | **Supplier name** | **Antibody** |
| --- | --- | --- |
| Ab3742 | Abcam | SUMO2/SUMO3 |
| sc5286 | Santa Cruz | αTubulin(B-7) |
| 926-68023 | Li COR | Donkey anti-Rabbit IgG |
| 926-32212 | Li COR | Donkey Anti-Mouse IgG |
| 100340 | BioLedgend | Anti Mouse CD3 |
| 102116 | BioLedgend | Anti Mouse CD28 |
| 300438 | BioLedgend | Anti Human CD3 |
| 377204 | BioLedgend | Anti Human CD28 |
| TS-5001-1C | MBL Life science | H-2Kb OVA Tetramer-SIINFEKL-PE |
| 127-005-099 | Jackson ImmunoResearch | AffiniPure® Goat Anti-Armenian Hamster IgG (H+L |
| 17090082 | Invitrogen | anti-mouse CD90.1 (Thy-1.1) -APC |
| 12090081 | Invitrogen | anti-mouse CD90.1 (Thy-1.1)-PE |
| 35589382 | Invitogen | anti-mouse KLRG1 PE-Cyanine5.5 |
| 363004 | Invitrogen | PE anti-human CD19 |
| 101320 | BioLegend | TruStain FcX™ (anti-mouse CD16/32) |
| 503808 | BioLegend | anti-mouse IL-2- PE |
| 104412 | BioLegend | anti-mouse CD62L - APC |
| 104450 | BioLegend | anti-mouse CD62L - APC/Fire™ 750 |
| 104543 | BioLegend | anti-mouse CD69- Brilliant Violet 785™ |
| IC8224G | RD systems | anti-mouse TCF7/TCF1 |
| 101916 | BioLegend | anti-mouse CD25 PE/Cyanine7 |
| 125224 | BioLegend | anti-mouse CD223 (LAG-3)- PE/Dazzle™ |
| 138429 | BioLegend | anti-mouse/human KLRG1- Brilliant Violet 785™ |
| 506341 | BioLegend | anti-mouse TNF-α- Brilliant Violet 785™ |
| 505840 | BioLegend | anti-mouse IFN-γ- Brilliant Violet 605™ |
| 100712 | BioLegend | anti-mouse CD8a-APC |
| 134010 | BioLegend | anti-mouse CD366 (Tim-3) PE/Cyanine7 |
| 103008 | BioLegend | anti-mouse/human CD44 - PE |
| 135043 | BioLegend | anti-mouse CD127 (IL-7Rα)- Brilliant Violet 650™ |
| 135231 | BioLegend | anti-mouse CD279 (PD-1) Brilliant Violet 711™ |
| 300326 | BioLegend | anti-human CD3- PerCP |
| 100725 | BioLegend | anti-mouse CD8a- Pacific Blue™ |
| 106318 | BioLegend | anti-mouse CTLA4 (CD152) PE/Dazzle™ 594 |
| 100222 | BioLegend | anti-mouse CD3- APC/Cyanine7 |

**Tables of primer sequences**

| **qPCR Primer sequence** | **Gene** | **Species** |
| --- | --- | --- |
| CAGGAGGCAAAACCTTCAAC :Forward | Sumo1 | Mouse |
| CTCCATTCCCAGTTCTTTCG :Reverse |  |  |
| ACGAGAAACCCAAGGAAGGA :Forward | Sumo2 |  |
| CTCCATTTCCAACTGTGCAG :Reverse |  |  |
| AGAAGCCCAAGGAGGGTGT :Forward | Sumo3 |  |
| CCTCGGGAGGCTGATCCT :Reverse |  |  |
| AGGTCGGTGTGAACGGATTTG :Forward | Gapdh |  |
| TGTAGACCATGTAGTTGAGGTCA :Reverse |  |  |
| TAGGCACCAGGGTGTGATG :Forward | Beta-Actin |  |
| GGGGTGTTGAAGGTCTCAAA :Reverse |  |  |
| GTGAATATATTAAACTCAAAGT :Forward | SUMO1 | Human |
| CTGACCCTCAAAGAGAAACC :Reverse |  |  |
| ATGGCCGACGAAAAGCCCAAG :Forward | SUMO2 |  |
| TGACAATCCCTGTCGTTCAC :Reverse |  |  |
| AGAATGACCACATCAACCTG :Forward | SUMO3 |  |
| CGAACCTGAATCTGATCTGC :Reverse |  |  |
| GGCCAACGAAAAGCCCACAGA :Forward | SUMO4 |  |
| GGTTCACAATAGGCTTTCATTAG :Reverse |  |  |
| ATG GAA ATC CCA TCA CCA TCTT :Forward | GAPDH |  |
| CGC CCC ACT TGA TTT TGG :Reverse |  |  |

| **PCR primer Sequence (Genotyping)** | **Gene** |
| --- | --- |
| CCATGGCGATTCTGAGATTCTGTCT : Forward | SUMO2 LoxP |
| AGTGGTGCCTCACAGGCCTTCAAT :Reverse |  |
| ATGGTGCCCAAGAAGAAGAG :Forward | LcKCre |
| CAGGTGCTGTTGGATGGTCT :Reverse |  |
| CAGCAGCAGGTGAGACAAAGT :Forward | OT1 |
| GGC TTT ATA ATT AGC TTG GTC C :Reverse |  |
| CAAATGTTGCTT GTCTGGTG :Forward | Internal positive control |
| GTCAGTCGAGTGCACAGTTT :Reverse |  |
